# Cholesterol-associated Locus *EHBP1* Protects against NASH Fibrosis

**DOI:** 10.1101/2023.07.08.548218

**Authors:** Fanglin Ma, Miriam Longo, Xin Huang, Marica Meroni, Satya Prakash, Erika Paolini, Dipankar Bhattacharya, Shuang Wang, Xiaobo Wang, Shama Mughal, Syed Hussain, Sumit Anand, Arif Yurdagul, Oren Rom, Paola Dongiovanni, Scott Friedman, Bishuang Cai

## Abstract

**SUMMARY:** Hepatic free cholesterol contributes to fibrosis in nonalcoholic steatohepatitis (NASH), but how hepatic cholesterol metabolism becomes dysregulated in NASH is not completely understood. We show here that human fibrotic NASH livers have decreased *EHBP1*, a novel GWAS locus associated with LDL-cholesterol, and that the *EHBP1* rs10496099 T>C variant in NASH patients is associated with decreased hepatic *EHBP1* expression and augmented NASH fibrosis. Congruent with the human data, EHBP1 loss- and gain-of-function increases and decreases NASH fibrosis in mice, respectively. Mechanistic studies reveal that EHBP1 promotes sortilin (SORT1)-mediated PCSK9 secretion, leading to LDLR degradation, decreased LDL uptake, and reduced TAZ, a fibrogenic effector. Moreover, we show that the TNFα/PPARα pathway suppresses EHBP1 in NASH. These data not only provide new mechanistic insight into the role of EHBP1 in cholesterol metabolism and NASH fibrosis by uncovering the interaction between EHBP1 and other cholesterol–related loci, including *SORT1*, *PCSK9*, and *LDLR*, but also elucidate a novel interplay between inflammation and EHBP1-mediated cholesterol metabolism.

*Highlights:* Hepatic EHBP1 is reduced in human and mouse fibrotic NASH *EHBP1* rs10496099 T>C is associated with decreased hepatic EHBP1 and augmented NASH EHBP1 reduces LDLR, cholesterol uptake, and TAZ in hepatocytes EHBP1 promotes sortilin-mediated PCSK9 secretion in hepatocytes TNFα suppresses EHBP1 expression in hepatocytes

## INTRODUCTION

Non-alcoholic fatty liver disease (NAFLD) is emerging as the leading cause of liver disease worldwide and is usually associated with heart disease, type 2 diabetes, and chronic kidney disease.^1–3^ NAFLD begins with the accumulation of large amounts of lipids in hepatocytes, and 20%–30% of subjects with NAFLD will develop non-alcoholic steatohepatitis (NASH), manifested by liver injury, inflammation, and fibrosis.^4^ NASH in turn can lead to the development of cirrhosis and hepatocellular carcinoma (HCC).^5^ Although several potential therapeutics are being tested in clinical trials, there are currently no FDA-approved drugs to treat NASH and its sequelae.^6, 7^ This is largely due to our lack of understanding of NASH progression to liver fibrosis, the most important predictor of long-term mortality, liver transplantation, and liver-related clinical events.^8^ Therefore, identifying novel factors or pathways that regulate liver fibrosis is critical for developing new therapeutic strategies for NASH.

NASH is likely driven by multiple factors, including insulin resistance, hepatosteatosis, and other insults that promote inflammation, fibrosis, and hepatocyte death.^9,10^ Emerging evidence shows that hepatic free cholesterol is an important mechanism to accelerate NASH progression.^11, 12^ Elevated hepatic cholesterol content is frequently seen in human NASH,^13, 14^ and high cholesterol diets have been found to promote NASH progression in mouse and nonhuman primate models.^15–19^ Hepatic cholesterol content can be regulated by proprotein convertase subtilisin/kexin type 9 (PCSK9), a secreted protein that binds to low-density lipoprotein receptor (LDLR) to induce its internalization and degradation in hepatocytes.^20–22^ PCSK9 inhibition has been shown to increase LDLR and the uptake of LDL cholesterol by the liver, leading to increased hepatic cholesterol, inflammation, and NASH fibrosis.^21, 23^ Cellular lipotoxicity caused by cholesterol accumulation can enhance hepatocyte death and activate liver macrophages and hepatic stellate cells (HSCs), leading to the development of progressive liver inflammation and fibrosis.^13^ We recently showed that cholesterol accumulation in hepatocytes blocks the proteasomal degradation of the Hippo effector TAZ, a key fibrosis driver in NASH.^24^ However, the regulation of cholesterol accumulation in hepatocytes during NASH is not completely understood.

Genome-wide association studies (GWAS) have revealed that several single nucleotide polymorphisms (SNPs) in *EHBP1*, the gene that encodes EH domain binding protein 1 (EHBP1), are associated with cholesterol.^25–28^ For instance, the intronic *EHBP1* SNP rs10496099 T>C variant is associated with decreased serum LDL and total cholesterol in a cohort of patients with coronary artery disease,^28^ though the mechanisms underlying this association and the role of EHBP1 in hepatic cholesterol metabolism are unknown. EHBP1 was originally identified as an interaction partner of the Eps15-homology domain-containing protein (EHD) family, which comprises four members (EHD1–4) essential for endocytic membrane trafficking. EHBP1 and EHDs have been implicated in regulating lipid metabolism in cultured cells,^29, 30^ but how they contribute to lipotoxicity-associated diseases including NASH is unknown. Published single-cell RNA sequencing (scRNA-seq) of human and mouse livers has revealed that hepatocytes express the highest level of EHBP1 among other liver cells and, surprisingly, hepatocyte EHBP1 expression is reduced in injured livers, including in cirrhotic livers with advanced fibrosis.^31, 32^ However, how EHBP1 may be mechanistically linked to cholesterol metabolism and NASH fibrosis remains unknown.

In this context, we now show that EHBP1 suppresses LDL receptor (LDLR) and LDL uptake in hepatocytes, leading to decreased TAZ and accordingly NASH fibrosis. Mechanistically, we demonstrate that EHBP1 promotes sortilin-mediated PCSK9 secretion by maintaining retromer homeostasis, leading to LDLR degradation. Moreover, in view of decreased EHBP1 expression in injured liver, we show that the NASH-relevant inflammatory cytokine TNFα suppresses EHBP1 expression by reducing the transcription factor peroxisome proliferator-activated receptor alpha (PPARα) in hepatocytes. These findings not only provide new mechanistic insight into the role of cholesterol-associated GWAS locus *EHBP1* in hepatic cholesterol metabolism and NASH fibrosis, but also elucidate a novel interplay between inflammation and EHBP1 mediated cholesterol metabolism in the context of NASH.

## RESULTS

### EHBP1 expression is reduced in human and mouse fibrotic NASH livers

Several independent GWAS have revealed that SNPs in *EHBP1* are associated with cholesterol metabolism.^25, 27, 33^ Given that the liver is the main organ involved in cholesterol metabolism and excessive hepatic cholesterol accelerates fibrotic NASH progression, we analyzed the *EHBP1* expression profile in liver cells. We interrogated a publicly available scRNA-seq dataset that examined liver cells isolated from human healthy livers versus cirrhotic livers with extensive fibrosis.^31^ We found that hepatocytes expressed the highest levels of *EHBP1* among all the liver cell types and that *EHBP1* expression was dramatically reduced in cirrhotic livers compared to healthy ones (**Figure S1A**). We then analyzed hepatocyte EHBP1 expression in human fibrotic NASH livers by co-staining EHBP1 and the hepatocyte marker HSA (Hepatocyte Specific Antigen) and found that, compared to the control group, hepatocyte EHBP1 was markedly reduced in NASH livers (**Figure 1A**). Consistently, EHBP1 expression was decreased in both human and nonhuman primate^18, 19^ fibrotic NASH livers (**Figures 1B, 1C, and S1B**). We also analyzed liver EHBP1 levels in several mouse models of NASH with fibrosis, including the Western diet combined with a low dose of carbon tetrachloride (WD+CCl_4_), the fructose, palmitate, cholesterol, and trans-fat diet (FPC), the high-fat, choline-deficient, L-amino-defined diet (CDAHFD), the high-fat, high-fructose, and high-cholesterol diet (FFC).^15, 16, 18, 34, 35^ Consistent with the human and nonhuman primate data, mRNA and protein levels of liver EHBP1 were significantly decreased in all the examined mouse NASH models (**Figures 1D, S1C, and S1D**). As the WD+CCl_4_ NASH model develops profound liver fibrosis in only 8–10 weeks and shares many features with human NASH,^15^ we used this model to examine the role of EHBP1 in liver fibrosis in NASH. We first performed unbiased RNA-seq from WD+CCl_4_–treated NASH livers and confirmed that *Ehbp1* expression was decreased in NASH livers (**Figure 1E**). This reduction was further confirmed by immunofluorescent staining of EHBP1 in hepatocytes (**Figure S1E**). Pathway ontology analysis revealed that processes involved in lipid, steroid, and cholesterol metabolism were significantly downregulated in mouse fibrotic NASH livers (**Figure 1F**). These human, nonhuman primate and mouse studies consistently show that EHBP1 is significantly suppressed in fibrotic NASH livers and prompt us to explore its causative role in this disease.

**Figure 1.**
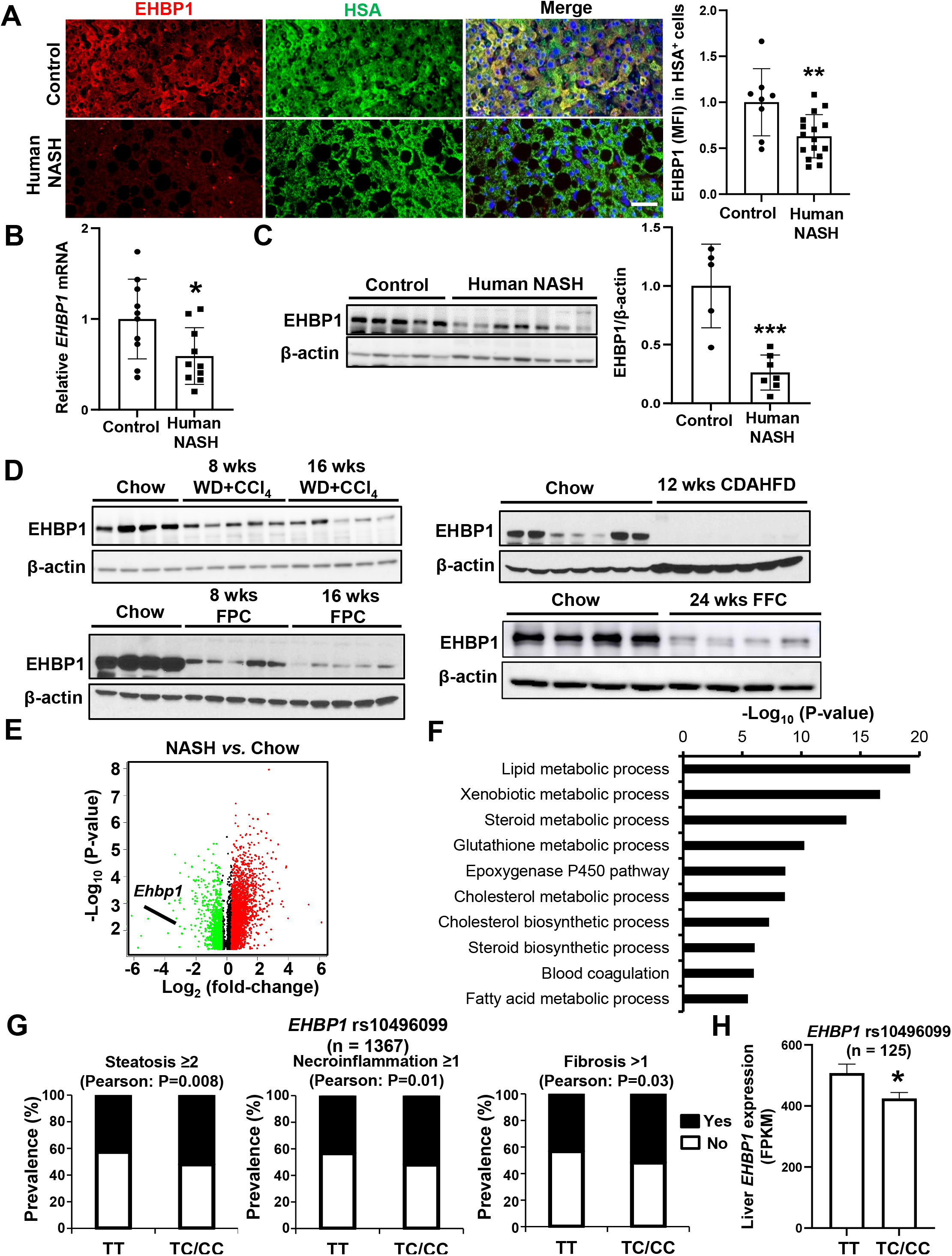
EHBP1 expression is reduced in human and mouse fibrotic NASH livers. (A) Immunofluorescent staining for hepatocyte marker HSA (green) and EHBP1 (red) and quantification of EHBP1 mean fluorescence intensity (MFI) in hepatocytes from control and NASH patients; DAPI counterstain for nuclei is shown (n = 8–16 subjects/group); Bar, 50 µm. (B) EHBP1 mRNA level in control and human NASH livers (n = 10 subjects/group). (C) Immunoblot of EHBP1 and densitometry quantification in control and human NASH livers, with β-actin as loading control. (D) Immunoblots of liver EHBP1 in NASH mice fed with different NASH diets (WD+CCl_4_, FPC, CDAHFD, and FFC) (n = 4–7 mice/group). (E) Volcano plot for the differentially expressed genes in NASH versus chow livers (n = 4 mice/group). (F) Gene ontology analysis for the downregulated genes as shown in panel E. (G) Pearson correlation analysis of the *EHBP1* rs10496099 C variant with liver damage in the Hepatology Service Cohort (n = 1367). (H) Liver *EHBP1* expression detected by RNA-seq from patients undergo bariatric surgery (*EHBP1* rs10496099 TT: n = 26; TC+CC: n = 99). FPKM, fragments per kilobase per million mapped reads. Data are presented as mean ± SEM. *p < 0.05, **p < 0.01, ***p < 0.001. See also Figure S1.

To determine whether cholesterol-associated *EHBP1* SNP rs10496099 T>C^28^ correlates with NASH progression, we assessed the *EHBP1* rs10496099 genotype in 1367 histologically characterized NAFLD patients from the Hepatology service cohort stratified according to disease severity. The clinical features of NAFLD patients stratified by the rs10496099 genotype and allele frequencies of the rs10496099 *EHBP1* T>C variant were shown in **Table S1** and **S2**. We found that the rs10496099 C allele was associated with a higher prevalence of steatosis ≥2, necroinflammation ≥1, and fibrosis >1 (**Figure 1G**). Importantly, a multivariate nominal regression analysis adjusted for sex, age, BMI (body mass index), and the *PNPLA3* G allele (the main genetic predictor of NAFLD) revealed that the rs10496099 C allele was associated with a higher risk of steatosis ≥2, necroinflammation ≥1, and fibrosis >1 (**Table S3**). Finally, we evaluated *EHBP1* expression in human liver biopsies (n = 125) from whom RNA-seq data were available^36^ (**Table S4**) and observed reduced mRNA levels in patients carrying the C allele (**Figure 1H**), indicating that the *EHBP1* rs10496099 T>C variant is loss-of-function. These human studies on the *EHBP1* SNP rs10496099 T>C variant indicate that EHBP1 may play a protective role against NASH progression.

### EHBP1 silencing in hepatocytes promotes liver fibrosis in NASH

We next explored a potential causative role of hepatocyte EHBP1 in NASH. As the whole-body EHBP1 knockout is embryonic lethal (unpublished data) and the EHBP1-floxed mouse line is currently not available, to explore the role of hepatocyte EHBP1 in NASH, we used an adeno associated virus type 8 (AAV8) vector with short hairpin RNA-targeting Ehbp1 (shEhbp1) driven by the H1 promoter (AAV8-H1-shEhbp1) to silence *Ehbp1* in hepatocytes.^16, 24^ We silenced EHBP1 by intravenous injection (IV) of AAV8-H1-shEhbp1 in C57BL/6J wild-type (WT) mice and examined liver histology, including fibrosis, steatosis, and inflammation after 10-week NASH diet (WD+CCl_4_)-feeding (**Figure 2A**). Despite similar body and liver weights (**Figure S2A**), we found that EHBP1 silencing induced liver fibrosis, as indicated by the increases in Sirius red staining, fibrotic gene expression, and αSMA^+^ area compared with the AAV8-H1-shCon (control)–treated group (**Figures 2B and S2B**). We also found that EHBP1 silencing increased liver steatosis and inflammation, as indicated by increased lipid droplet area, and increased inflammatory cells and inflammatory genes—*Adgre1, Tnfa* and *Ccl2*, and %F4/80^+^ area (**Figures 2C, 2D, and S2C**). Consistent with the liver histology feature, plasma ALT and AST levels were enhanced in shEhbp1-treated NASH mice (**Figure 2E**), indicating that liver injury was elevated in hepatocyte EHBP1–silenced NASH mice. These *in vivo* studies uncover that the silencing of hepatocyte EHBP1 promotes NASH by accelerating liver fibrosis, steatosis, and inflammation.

**Figure 2.**
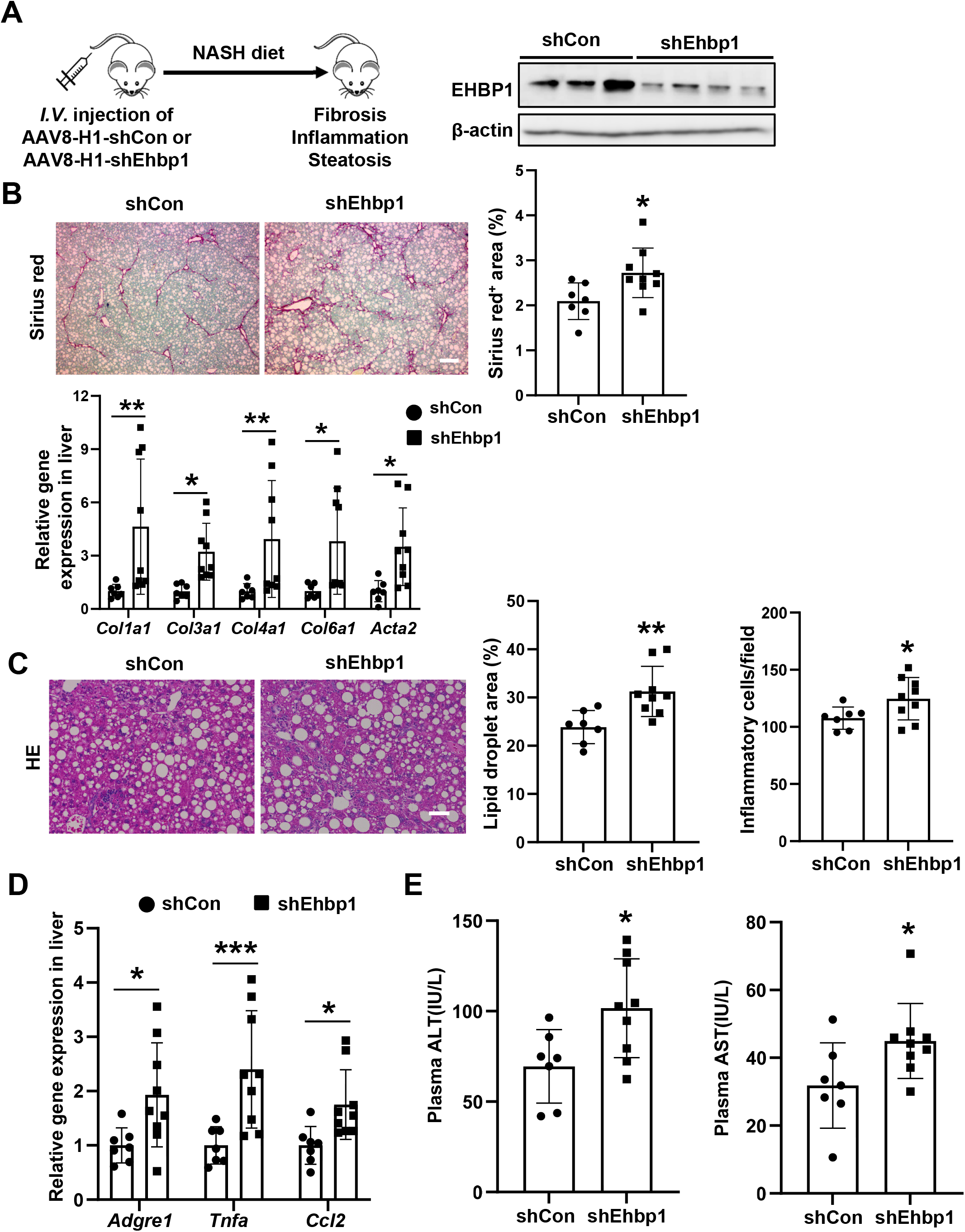
EHBP1 silencing in hepatocytes promotes liver fibrosis in NASH. The following parameters were measured in male C57BL/6J mice treated with AAV8-H1-shEhbp1 or AAV8-H1-shCon (control) virus and then fed NASH diet (WD+CCl_4_) for 10 weeks. (A) Immunoblot of liver EHBP1, with β-actin as loading control. (B) Staining of liver sections for Sirius red and quantification of percent Sirius red^+^ area by Image J. Liver mRNA levels of *Col1a1*, *Col3a1*, *Col4a1*, *Col6a1*, and *Acta2*; Bar, 200 µm. (C) Staining of liver sections for H&E and quantification of lipid droplet area and inflammatory cells by Image J; Bar, 50 µm. (D) Liver mRNA levels of *Adgre1* (F4/80), *Tnfa*, and *Ccl2*. (E) Plasma ALT and AST. Data are presented as mean ± SEM. *p < 0.05, **p < 0.01, ***p < 0.001 (n = 7–9 mice/group). See also Figure S2.

### EHBP1 overexpression in hepatocytes eliminates liver fibrosis in NASH

To explore a potential protective role of hepatocyte EHBP1 against NASH, we overexpressed EHBP1 specifically in hepatocytes by using TBGS1 (shortened version of the liver-specific thyroxine-binding globulin (TBG) promoter).^37^ We intravenously injected AAV8-TBGS1-eGFP (control) or AAV8-TBGS1-Ehbp1 in WT mice and placed them on a NASH diet (WD+CCl_4_) for 10 weeks (**Figure 3A**). At the endpoint, the two groups of mice had similar body and liver weights (**Figure S3A**). However, in sharp contrast to EHBP1 silencing, EHBP1 overexpression in hepatocytes reduced liver fibrosis, as indicated by the decreases in Sirius red staining, fibrotic gene expression, and the αSMA^+^ area in AAV8-TBGS1-Ehbp1-treated NASH mice (**Figures 3B and S3B**). Consistently, this was accompanied by decreases in steatosis, liver inflammation, and plasma ALT and AST (**Figures 3C–3E and S3C**). Taken together, our *in vivo* EHBP1 loss- and gain-of-function studies demonstrate that hepatocyte EHBP1 plays a protective role against NASH.

**Figure 3.**
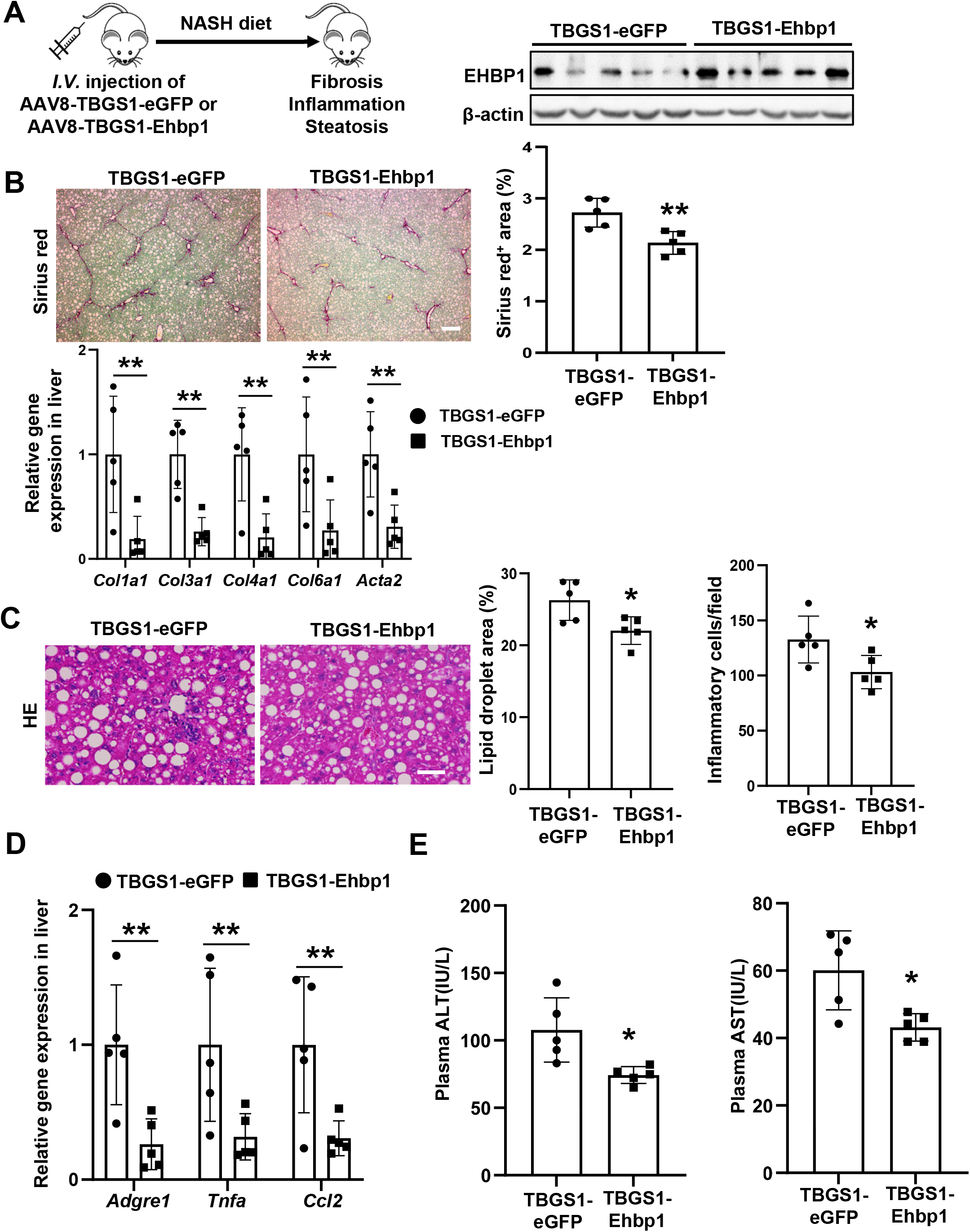
EHBP1 overexpression in hepatocytes eliminates liver fibrosis in NASH. The following parameters were measured in male C57BL/6J mice treated with AAV8-TBGS1-eGFP (control) or AAV8-TBGS1-Ehbp1 virus and then fed NASH diet (WD+CCl_4_) for 10 weeks. (A) Immunoblot of liver EHBP1, with β-actin as loading control. (B) Staining of liver sections for Sirius red and quantification of percent Sirius red^+^ area by Image J. Liver mRNA levels of *Col1a1*, *Col3a1, Col4a1*, *Col6a1,* and *Acta2*; Bar, 200 µm. (C) Staining of liver sections for H&E and quantification of lipid droplet area and inflammatory cells by Image J; Bar, 50 µm. (D) Liver mRNA levels of *Adgre1 (F4/80), Tnfa*, and *Ccl2*. (E) Plasma ALT and AST. Data are presented as mean ± SEM. *p < 0.05, **p < 0.01 (n = 5 mice/group). See also Figure S3.

### EHBP1 suppresses hepatic LDLR, free cholesterol, and TAZ in NASH

Given that *EHBP1* SNPs are associated with LDL cholesterol,^25, 26, 38, 39^ we sought to determine whether EHBP1 regulates LDLR, a receptor mediating LDL cholesterol uptake into hepatocytes.^40^ Interestingly, while *Ldlr* mRNA was not altered in livers from mice treated with AAV8-H1-shEhbp1, LDLR protein levels were significantly increased (**Figures 4A and S4A**), which was associated with elevated free cholesterol, as indicated by the enhanced Filipin staining, a probe for free cholesterol^24^ (**Figure 4B**). In contrast to hepatocyte EHBP1 silencing result, EHBP1 overexpression resulted in significantly reduced LDLR protein and free cholesterol levels without altering *Ldlr* mRNA (**Figures 4C, 4D, and S4B**). To determine whether EHBP1 directly regulates LDLR in hepatocytes, we silenced EHBP1 in isolated primary mouse hepatocytes by using two different siRNAs. As found in the *in vivo* studies, EHBP1 silencing in primary hepatocytes induced the protein level of LDLR but not its mRNA level (**Figures 4E and S4C**). Consistent with the increased LDLR, the uptake of LDL was enhanced in EHBP1-silenced hepatocytes (**Figure 4F**). Our recent study showed that cholesterol prevented the proteasomal degradation of TAZ, a Hippo effector that acted as a transcription factor to induce Indian Hedgehog (IHH) expression in hepatocytes, leading to HSC activation and liver fibrosis in NASH.^16, 24^ Consistent with this concept, we found that cholesterol delivered by LDL particles that contain approximately 50% cholesterol^41^ markedly increased TAZ in primary hepatocytes (**Figure 4G**). We next determined whether EHBP1 regulates TAZ stability. As expected, hepatocyte EHBP1–silenced NASH livers had increased TAZ at the protein level (**Figure 4H**) and increased *Ihh* mRNA (**Figure S4D**). Of note, EHBP1 silencing did not significantly change *Wwtr1* (TAZ) mRNA and the protein level of YAP, another Hippo effector (**Figures S4E and 4H**). In contrast to EHBP1 silencing, EHBP1 overexpression reduced TAZ protein levels and suppressed *Ihh* mRNA (**Figures 4I, S4F, and S4G**). To further confirm the link between EHBP1 and TAZ, we knocked down EHBP1 in isolated primary hepatocytes. Consistent with the *in vivo* data, we found that EHBP1 silencing enhanced TAZ protein levels in primary mouse hepatocytes (**Figures 4J and S4H**). The increased TAZ was also confirmed with the mouse hepatocyte cell line AML12 upon EHBP1 silencing (**Figure S4I**). These data indicate that EHBP1 protects against NASH fibrosis by reducing hepatic cholesterol and TAZ.

**Figure 4.**
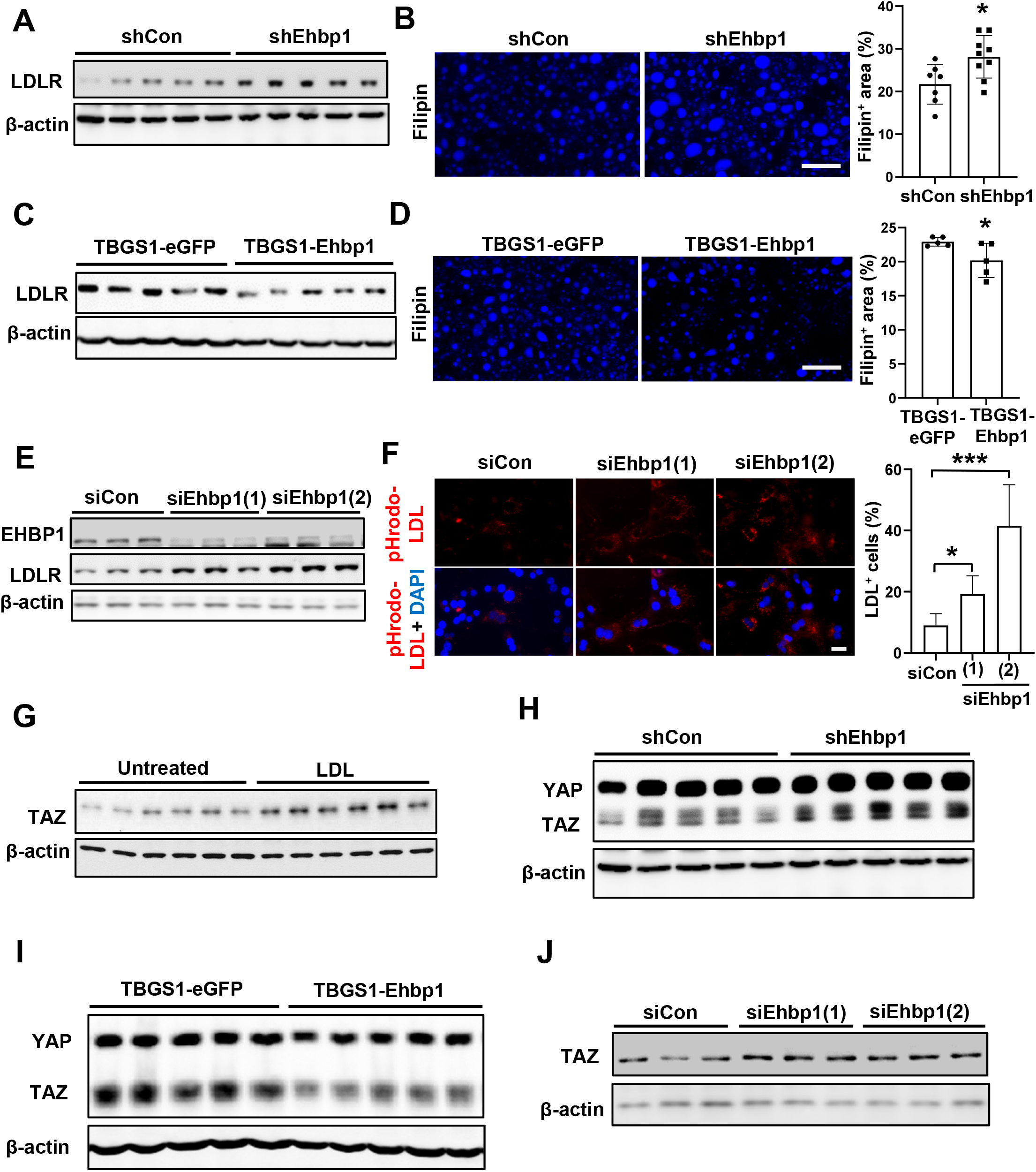
EHBP1 suppresses hepatic LDLR, cholesterol, and TAZ in NASH. (A) Immunoblot of liver LDLR from NASH mice treated with AAV8-H1-shCon or AAV8-H1-shEhbp1. (B) Filipin staining of NASH liver sections and quantification from NASH mice treated with AAV8-H1-shCon or AAV8-H1-shEhbp1 (n = 7–9 mice/group); Bar, 50 µm. (C) Immunoblot of liver LDLR from NASH mice treated with AAV8-TBGS1-eGFP or AAV8-TBGS1-Ehbp1. (D) Filipin staining of NASH liver sections and quantification from NASH mice treated with AAV8-TBGS1-eGFP or AAV8-TBGS1-Ehbp1 (n = 5 mice/group); Bar, 50 µm. (E) Immunoblots of EHBP1 and LDLR in primary mouse hepatocytes transfected with siCon or siEhbp1. (F) pHrodo-LDL uptake with quantification in primary mouse hepatocytes transfected with siCon or siEhbp1 (n = 3 biological replicates); Bar, 100 µm. (G) Immunoblot of TAZ in primary mouse hepatocytes that were incubated with 100 µg/ml LDL for 8 h. (H) Immunoblots of YAP and TAZ in AAV8-H1-shCon or AAV8-H1-shEhbp1-treated NASH livers. (I) Immunoblots of YAP and TAZ in AAV8-TBGS1-eGFP or AAV8-TBGS1-Ehbp1-treated NASH livers. (J) Immunoblot of TAZ in primary mouse hepatocytes transfected with siCon or siEhbp1. Data are presented as mean ± SEM. *p < 0.05, ***p < 0.001. See also Figure S4.

### EHBP1 promotes sortilin-mediated PCSK9 secretion

We next sought to determine the mechanism by which EHBP1 reduces LDLR in hepatocytes. Given that our *in vivo* and *in vitro* studies showed that EHBP1 did not alter *Ldlr* mRNA, we examined whether EHBP1 plays a role in PCSK9-mediated LDLR degradation (**Figure S5A**). As EHBP1 has been shown to promote the internalization of cell-surface proteins (*e.g.,* transferrin receptor and GLUT4) and PCSK9-induced LDLR internalization is required for LDLR degradation,^30, 42–44^ we first examined whether EHBP1 silencing abolishes the ability of PCSK9 to degrade LDLR. However, we found that EHBP1 was not required for PCSK9-mediated LDLR degradation (**Figure S5B**), indicating that EHBP1 does not affect PCSK9-induced LDLR internalization and degradation. We then explored whether EHBP1 regulates PCSK9 secretion. Surprisingly, we found that plasma PCSK9 levels were decreased in EHBP1-silenced NASH mice and increased in EHBP1-overexpressed NASH mice (**Figure 5A**). Consistent with the concept that PCSK9-mediated LDLR degradation leads to hypercholesterolemia,^45^ we found that decreased plasma PCSK9 was associated with decreased plasma LDL and total cholesterol in EHBP1-silenced NASH mice, while increased plasma PCSK9 was associated with increased plasma LDL and total cholesterol in EHBP1-overexpressed NASH mice (**Figures 5B and 5C**). Interestingly, a recent study showed that the *EHBP1* rs10496099 C allele is associated with decreased serum LDL and total cholesterol.^28^ As hepatic *EHBP1* expression is lower in NAFLD patients carrying the rs10496099 C allele (**Figure 1H**), reduced hepatic EHBP1 expression in humans is associated with decreased circulating cholesterol, which is consistent with our EHBP1-silenced NASH mouse studies. Given that *Pcsk9* mRNA was not altered in either EHBP1-silenced or EHBP1-overexpressed NASH livers (**Figure S5C**), EHBP1-induced plasma PCSK9 is not regulated via modulating *Pcsk9* expression. We then assayed whether EHBP1 regulates PCSK9 secretion in primary hepatocytes and found that EHBP1 silencing reduced secreted PCSK9 in the culture media from both primary mouse and human hepatocytes without changing *Pcsk9* mRNA in hepatocytes (**Figures 5D, 5E, S5D, and S5E**), which suggests that EHBP1 promotes PCSK9 secretion. Since PCSK9 secretion from hepatocytes depends on sortilin, encoded by the hypercholesterolemia-risk gene *SORT1*,^46^ we speculated that EHBP1 facilitates PCSK9 secretion by regulating sortilin. Indeed, we found that sortilin was decreased in EHBP1-deficient primary mouse and human hepatocytes (**Figures 5F and 5G**). The decreased sortilin and PCSK9 secretion were also confirmed in EHBP1-silenced AML12 cells (**Figures S5F and S5G**). Lastly, we found that hepatocyte sortilin was decreased in both human and mouse NASH livers where EHBP1 expression was reduced (**Figures 5H and S5H**). These data suggest that EHBP1 promotes PCSK9 secretion by enhancing sortilin.

**Figure 5.**
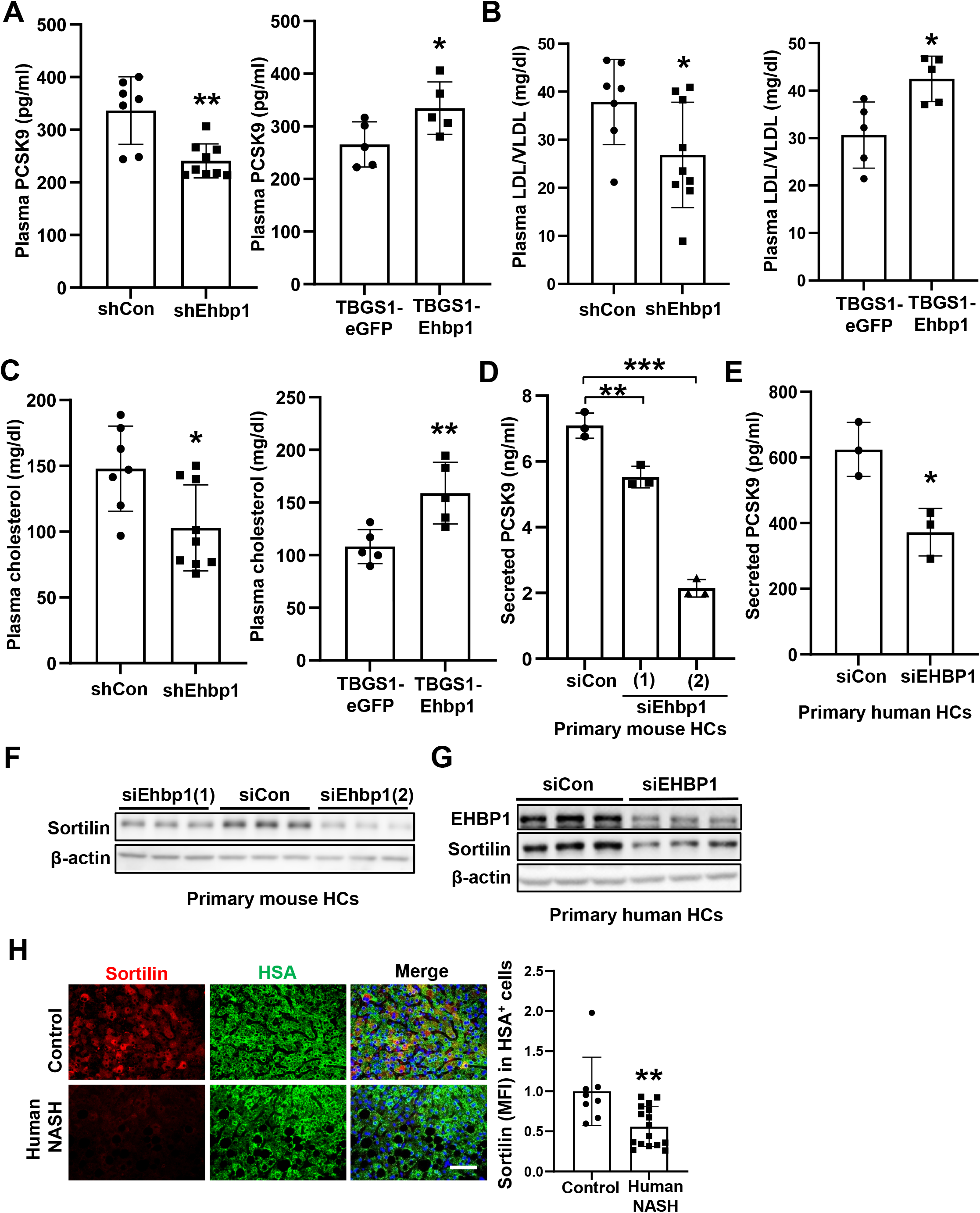
EHBP1 promotes sortilin-mediated PCSK9 secretion. (A) Plasma PCSK9 levels in NASH mice (n = 5–9 mice/group). (B) Plasma LDL/VLDL cholesterol levels in NASH mice (n = 5–9 mice/group). (C) Plasma total cholesterol levels in NASH mice (n = 5–9 mice/group). (D) Secreted PCSK9 levels were measured in culture media from primary mouse hepatocytes treated with siCon or siEhbp1 (n = 3 biological replicates). (E) Secreted PCSK9 levels were measured in culture media from primary human hepatocytes treated with siCon or siEHBP1 (n = 3 biological replicates). (F) Immunoblot of sortilin in primary mouse hepatocytes treated with siCon or siEhbp1 (n = 3 biological replicates). (G) Immunoblots of EHBP1 and sortilin in primary human hepatocytes treated with siCon or siEHBP1 (n = 3 biological replicates). (H) Immunofluorescent staining for HSA (green) and sortilin1 (red) and quantification of sortilin MFI in HSA^+^ cells from control and NASH patients; DAPI counterstain for nuclei is shown (n = 8–16 subjects/group); Bar, 50 µm. Data are presented as mean ± SEM. *p < 0.05, **p < 0.01, ***p < 0.001; HC, hepatocyte. See also Figure S5.

### EHBP1 stabilizes sortilin by maintaining retromer homeostasis

We next explored the potential mechanisms by which EHBP1 induces sortilin. As *Sort1* mRNA levels were not significantly changed in either EHBP1-silenced or EHBP1-overexpressed NASH livers (**Figure S6A**), we sought to determine whether EHBP1 promotes sortilin stability at the protein level. Emerging evidence suggests that sortilin undergoes retrograde transport from endosomes to the *trans*-Golgi network (TGN) by the retromer complex, which contains two subcomplexes: a vacuolar protein sorting-associated protein 26 (Vps26)-Vps29-Vps35 trimeric subcomplex and a membrane-associated sorting nexin (SNX) dimer.^47^ Retromer-mediated retrograde transport of sortilin can stabilize sortilin through escaping from its lysosomal degradation.^48–50^ To determine whether EHBP1 interacts with sortilin and the retromer, we performed co-immunoprecipitation (Co-IP) experiments and revealed that EHBP1 interacted with sortilin and retromer units—Vps35 and Vps26a in isolated primary hepatocytes and AML12 cells (**Figures 6A and S6B**). Of note, EHD1, a known EHBP1 binding partner, served as a positive control for the Co-IP assay. Confocal microscopy confirmed that EHBP1 partially co-localized with sortilin and Vps35 retromer in hepatocytes (**Figures 6B and S6C**). To further establish the link between sortilin and retromer function in hepatocytes, we knocked down retromer units, including Vps26a, Vps35 and SNX1, and determined the effect of retromer on sortilin. As expected, the deficiency of the retromer complex dramatically reduced sortilin in primary hepatocytes and AML12 cells (**Figures 6C and S6D**). Of note, Vps26a silencing reduced Vps35, and vice versa. This reciprocal effect of Vps26a and Vps35 was consistent with previous results reported in HeLa cells.^51^ We next evaluated the importance of retromer mediated sortilin stability in our described PCSK9 secretion-LDLR-TAZ pathway. Consistent with decreased sortilin in retromer-deleted hepatocytes, we found that the deficiency of the retromer complex reduced PCSK9 secretion from primary hepatocytes, which was associated with increased LDLR and TAZ (**Figures 6D and 6E**). As our data indicate that retromer stabilizes sortilin and EHBP1 interacts with the retromer complex in hepatocytes, we next sought to determine whether EHBP1 stabilizes sortilin by regulating intracellular retromer. Although the role of EHBP1 in regulating retromer function is unknown, its binding partner EHD1 has been shown to facilitate retromer-mediated retrograde transport by controlling the intracellular localization of retromer in HeLa cells.^51, 52^ For instance, EHD1-depletion altered subcellular localization of Vps26a from the cell periphery to the perinuclear region, leading to a defect in the retrograde transport.^52^ Thus, we speculated that EHBP1 might alter the subcellular localization of Vps26a in hepatocytes. Interestingly, we found that, similar to EHD1 deficiency, EHBP1 silenced hepatocytes had increased perinuclear Vps26a (**Figure 6F**). Thus, EHBP1 may stabilize sortilin by controlling the intracellular localization and homeostasis of the retromer.

**Figure 6.**
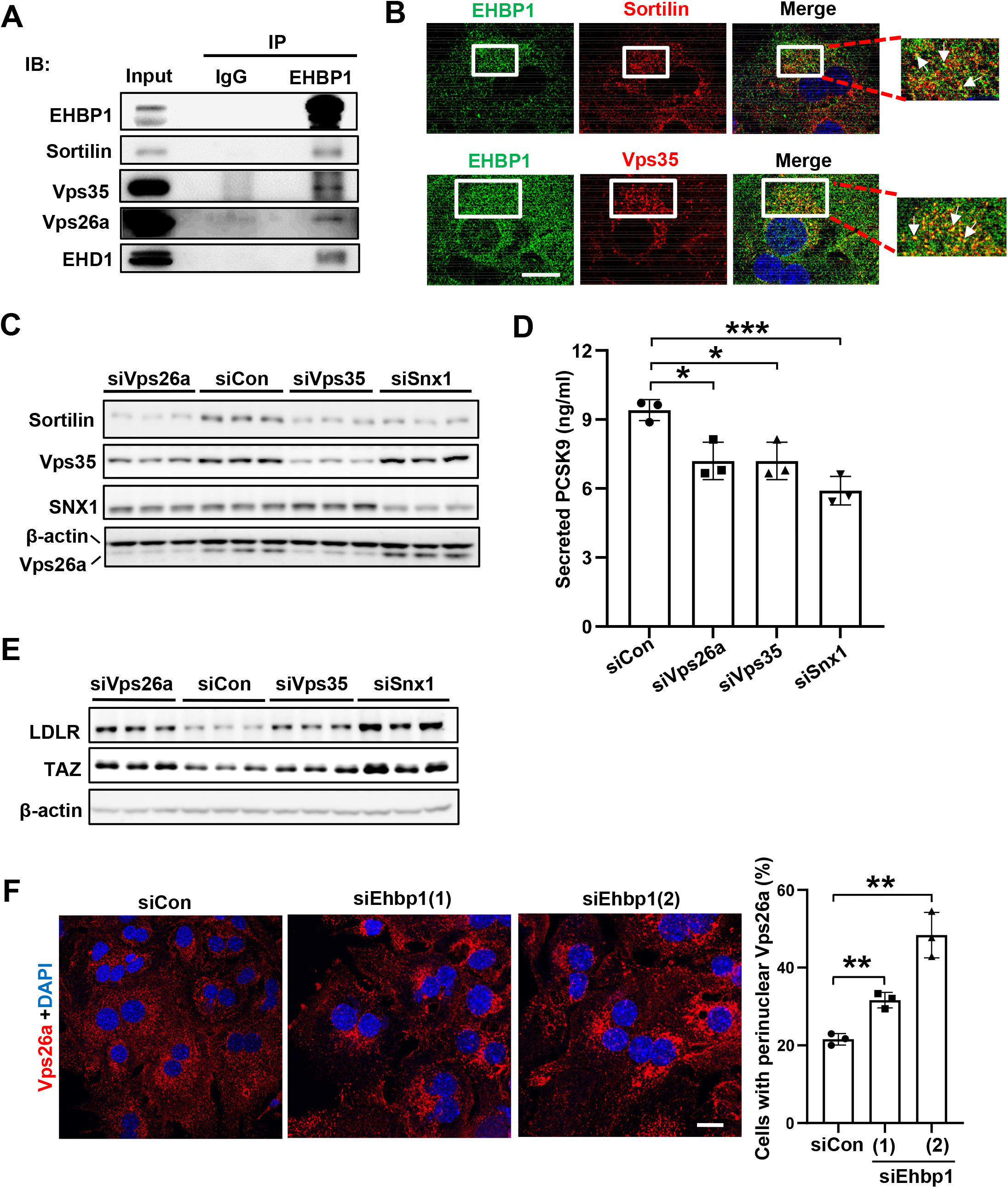
EHBP1 stabilizes sortilin by maintaining retromer homeostasis. (A) Co-immunoprecipitation of EHBP1, sortilin, Vps35, Vps26a, and EHD1 in primary mouse hepatocytes. (B) Confocal imaging of EHBP1, sortilin, and Vps35 in primary mouse hepatocytes; DAPI counterstain for nuclei is shown and white arrows indicate colocalization; Bar, 20 µm. (C) Immunoblots of sortilin, Vps35, SNX1, and Vps26a in primary mouse hepatocytes treated with siRNAs. (D) Secreted PCSK9 levels were measured in culture media from primary mouse hepatocytes treated with siRNAs (n = 3 biological replicates). (E) Immunoblots of LDLR and TAZ in primary mouse hepatocytes treated with siRNAs. (F) Confocal imaging of Vps26a in primary mouse hepatocytes treated with siCon or siEhbp1; DAPI counterstain for nuclei and quantification of percent cells with perinuclear Vps26a are shown (n = 3 biological replicates); Bar, 20 µm. Data are presented as mean ± SEM. *p < 0.05, **p < 0.01, ***p < 0.001. See also Figure S6.

### TNFα suppresses EHBP1 expression in hepatocytes

Increased inflammation is a hallmark of NASH and plays a pivotal role in NASH progression through upregulating key molecules associated with lipid metabolism and fibrosis in the liver. As previous studies showed that inflammatory cytokines, including TNFα, induced LDLR and LDL cholesterol accumulation in hepatocytes^53–55^ and our new data showed that EHBP1 expression was suppressed in NASH (**Figure 1**), we next examined the potential role of increased TNFα as a suppressor of EHBP1 in NASH. To study whether TNFα regulates EHBP1, we treated primary mouse hepatocytes with recombinant TNFα and found that TNFα treatment reduced EHBP1 and increased TAZ (**Figures 7A and S7A**), suggesting that suppressed EHBP1 during NASH (**Figure 1**) may result from increased TNFα (**Figure 7B**). We next explored the mechanism by which TNFα reduces EHBP1. We searched the PathwayNet database for potential transcription factors that control EHBP1 and found that PPARα is one of the top candidates. Interestingly, TNFα treatment suppressed the expression of both *Ppara* and *Ehbp1* (**Figure 7C**). To determine whether PPARα directly binds to the promoter region of *Ehbp1*, we analyzed a published chromatin immunoprecipitation-sequencing (ChIP-seq) of PPARα from WT versus *Ppara^-/-^* livers.^56^ As expected, we observed that PPARα activation with agonist enhanced the binding of PPARα to the *Ehbp1* promoter region in WT livers, which was abolished in *Ppara^-/-^* livers (**Figure 7D**), indicating that PPARα may serve as a transcription factor for EHBP1. In line, when *Ppara* was silenced using siRNA, we found that PPARα deficiency decreased EHBP1 expression in primary hepatocytes (**Figure 7E**). The reduced EHBP1 was also confirmed in primary hepatocytes isolated from mice with hepatocyte-specific deletion of *Ppara* (*Ppara^HKO^*) compared to *Ppara^fl/fl^* (**Figure S7B**). Similar to EHBP1, *PPARA* expression in hepatocytes was dramatically reduced in human cirrhotic livers and *PPARA* expression was also found to decrease in nonhuman primate NASH livers as well as in FPC- and FFC-induced mouse NASH livers (**Figures S7C–S7F**). Moreover, previous studies showed that the activation of PPARα is suppressed in FPC- and CDAHFD-induced NASH livers and that hepatocyte PPARα deficiency increased hepatic cholesterol and augmented NASH.^57–59^ Along this line, analysis of single nucleus RNA sequencing (snRNA-seq) of livers from our WD+CCl_4_- NASH mouse model^32^ revealed that both *Ehbp1* and *Ppara* were downregulated in hepatocytes (**Figure 7F**). Finally, in line with decreased EHBP1, PPARα silencing in primary mouse hepatocytes also led to reduced sortilin and increased LDLR and TAZ (**Figure 7G**). Together, these data indicate that TNFα suppresses EHBP1 expression by suppressing PPARα in hepatocytes during NASH progression.

**Figure 7.**
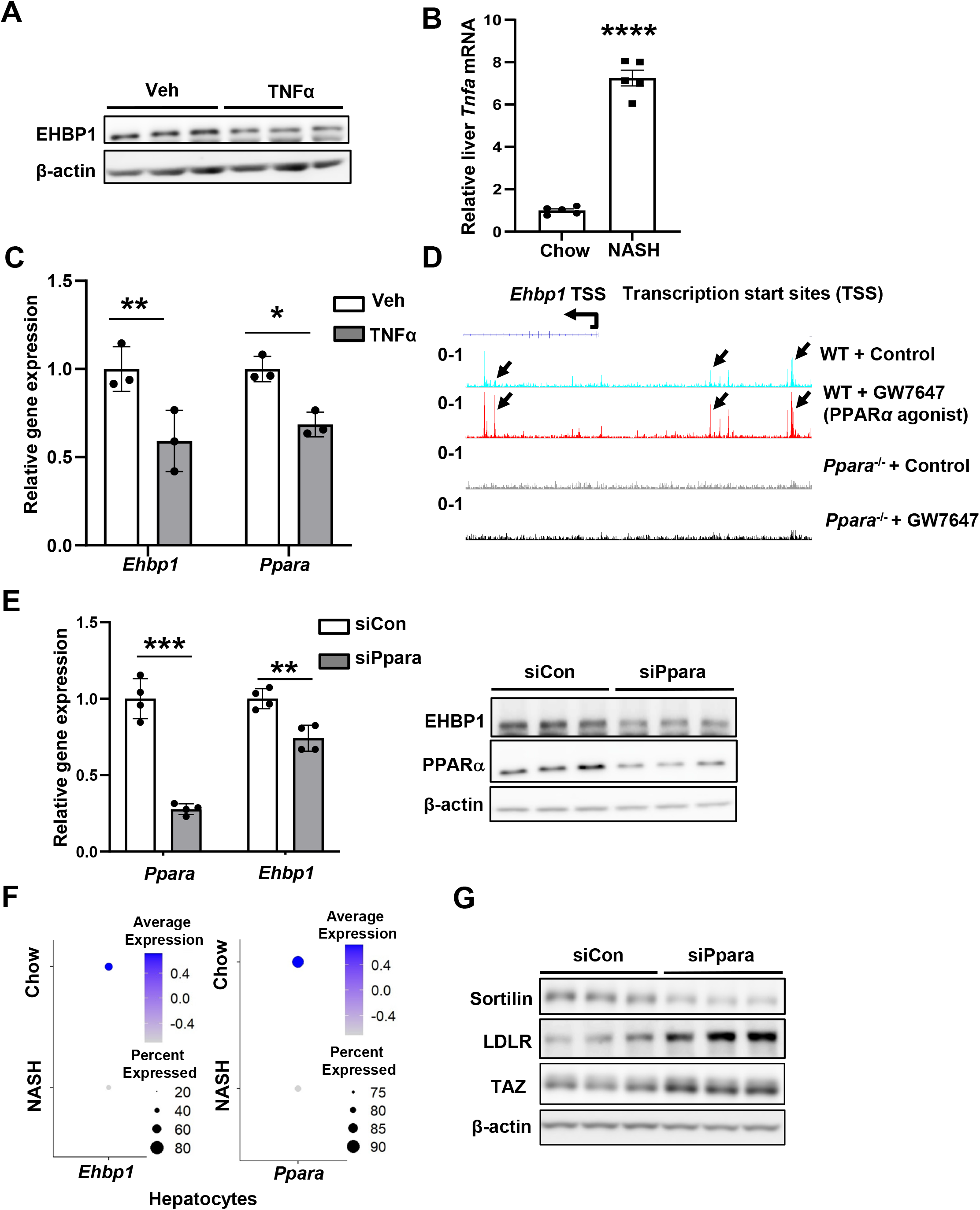
TNFα suppresses EHBP1 expression in hepatocytes. (A) Immunoblot of EHBP1 in primary mouse hepatocytes that were incubated with 20 ng/ml TNFα for 24 h. (B) *Tnfa* mRNA level in NASH (WD+CCl_4_) livers (n = 5 mice/group). (C) mRNA levels of *Ehbp1* and *Ppara* in primary mouse hepatocytes that were incubated with 20 ng/ml TNFα for 24 h (n = 3 biological replicates). (D) Genome browser tracks of PPARα ChIP-seq at *Ehbp1* promoter region in GW7647-treated WT or *Ppara^-/-^*livers. (E) mRNA and protein levels of EHBP1 and PPARα in primary mouse hepatocytes treated with siCon or siPpara (n = 3 biological replicates). (F) Dot plot for the expression of *Ehbp1* and *Ppara* in hepatocytes from single nuclear RNA-seq of chow and NASH livers. Average gene expression and percentage of cells that express each gene are presented with differential color intensities and circle sizes, respectively. (G) Immunoblots of sortilin, LDLR, and TAZ in primary mouse hepatocytes treated with siCon or siPpara. Data are presented as mean ± SEM. *p < 0.05, **p < 0.01, ***p < 0.001, ****p < 0.0001. See also Figure S71.

## DISCUSSION

The primary finding in this study is the discovery of a novel mechanism by which cholesterol associated GWAS locus *EHBP1* protects against NASH fibrosis. The protective role of EHBP1 is supported by the evidence that the loss-of-function *EHBP1* rs10496099 T>C genetic variation in NAFLD patients is associated with more liver fibrosis. Consistent with the human study, EHBP1-silenced and EHBP1-overexpressed NASH mice have increased and decreased liver fibrosis, respectively. Our mechanistic studies demonstrate that EHBP1 facilitates sortilin mediated PCSK9 secretion and decreases LDLR and cholesterol uptake in hepatocytes, leading to the suppression of hepatocyte TAZ and NASH fibrosis (**Figure S7G**). However, the protective effect of EHBP1 is abolished by the TNFα-mediated suppression of EHBP1 expression during NASH. Our study reveals a novel link between EHBP1 and sortilin-mediated PCSK9 secretion in hepatocytes. Moreover, at a cellular level, we show that EHBP1 stabilizes sortilin by maintaining retromer homeostasis. These findings provide new mechanistic insight into the regulation of cholesterol metabolism in hepatocytes and reveal a novel interplay between inflammation and cholesterol metabolism in NASH.

Our finding that EHBP1 promotes PCSK9 secretion by maintaining retromer homeostasis and stabilizing sortilin is novel, as the roles of EHBP1 and retromer in PCSK9 secretion have never been described before. Moreover, we elucidate a novel link among the four cholesterol associated loci *EHBP1*, *SORT1*, *PCSK9,* and *LDLR*. Sortilin has been shown to co-localize with PCSK9 in the TGN and facilitate its secretion from hepatocytes.^46^ Similar to our hepatocyte EHBP1–silenced mice, hepatocyte-specific sortilin knockout mice have decreased plasma PCSK9, increased liver LDLR, and reduced plasma LDL cholesterol.^46, 60^ In addition to facilitating PCSK9 secretion, hepatocyte sortilin has been shown to modulate very-low-density lipoprotein (VLDL) secretion,^60–64^ therefore, whether EHBP1 regulates VLDL secretion by stabilizing sortilin needs further investigation. Studies in adipocytes and HepG2 cells showed that sortilin can be recycled by retromer-mediated retrograde transport from endosomes back to the TGN, avoiding degradation in lysosomes.^48, 49^ Consistent with this concept, we demonstrated that retromer subunits including Vps26a, Vps35, and SNX1 stabilize sortilin and promote PCSK9 secretion in primary hepatocytes. Future *in vivo* studies will be required to investigate the roles of sortilin and the retromer complex in NASH fibrosis. Another important question that requires further investigation is how EHBP1 regulates the retromer function in hepatocytes. In a state of homeostasis, the retromer complex is localized to endosomes and facilitates the export of endosomal cargo to the TGN via retrograde transport.^65, 66^ The binding partner of EHBP1, EHD1, has been shown to interact with retromer on endosomes and promotes efficient retrograde transport in HeLa cells.^51, 52^ These studies showed that EHD1 deletion or mutation causes redistribution of retromer from peripheral endosomes to the perinuclear region and abolishes retromer-mediated retrograde transport.^51, 52^ Consistent with this concept, we demonstrate that EHBP1, EHD1, and retromer form a protein complex in primary hepatocytes and that, like EHD1, EHBP1 deficiency favors a perinuclear localization of retromer. Thus, it is likely that EHBP1 cooperates with EHD1 to maintain retromer homeostasis and facilitate efficient retromer mediated retrograde transport of sortilin. As EHBP1 binds to the EH domain in EHD1 through its Asn-Pro-Phe (NPF) motifs,^30, 67^ It will be interesting to determine whether blocking the interaction between EHBP1 and EHD1 by mutating the NPF motifs or deleting the EH domain disrupts retromer homeostasis and suppresses the retrograde transport of sortilin.

LDLR-mediated LDL uptake is a major cholesterol input pathway for the liver, and our data indicate that hepatocyte EHBP1 blocks cholesterol uptake, which in turn decreases TAZ and liver fibrosis in NASH. Although our study focuses on the anti-fibrotic effect of hepatocyte EHBP1, we also show that EHBP1 protects against steatosis and inflammation in NASH. As LDL particles contain 50% cholesterol and 5% triglyceride,^68, 69^ EHBP1-reduced steatosis may result from decreased LDL uptake. In view of the anti-inflammatory effect of EHBP1, since cholesterol accumulation can induce inflammation,^70, 71^ it is likely that EHBP1-suppressed inflammation is simply due to the decreased hepatic cholesterol contents. Another possibility is through decreased TAZ, seen in EHBP1-overexpressed NASH livers, as we previously showed that hepatocyte TAZ has a pro-inflammatory role in NASH.^16^ Our investigation into why EHBP1 expression is reduced in NASH livers led to the discovery of a positive feedback loop between cholesterol accumulation and inflammation. On the one hand, excessive cholesterol can directly induce inflammatory pathways.^71^ On the other hand, inflammatory signaling can disrupt cholesterol homeostasis and exacerbate cholesterol accumulation.^54, 72^ Several studies have focused on the effects of cholesterol on inflammatory pathways, but only a few have investigated the effects of inflammation on cholesterol metabolism. We now show that NASH-relevant inflammatory cytokine TNFα suppresses EHBP1 expression and increases TAZ. In line with our finding, TNFα has been shown to enhance LDL uptake and exacerbate cholesterol accumulation by increasing LDLR in fatty livers and hepatocytes.^53, 55, 73^ Therefore, our study provides evidence that inflammation can amplify cholesterol dysregulation, such as via abolishing EHBP1-mediated cholesterol homeostasis. The maintenance of hepatic cholesterol homeostasis requires a balance between cholesterol input and output.^74, 75^ Although our study focuses on the liver cholesterol input pathway, whether EHBP1 plays a role in regulating liver cholesterol output, *e.g.*, catabolizing cholesterol to bile acids is unknown. Given that bile acid signaling via farnesoid X receptor (FXR) also regulates NASH progression and cholesterol metabolism,^76^ whether EHBP1 modulates bile acid signaling requires further investigation.

Dyslipidemia is a common risk factor for both atherosclerotic cardiovascular disease (ASCVD) and NASH.^77^ However, lowering circulating LDL cholesterol with statins has shown inconsistent results in improving NASH outcomes despite potent effect in ameliorating the risk of ASCVD.^78^ Statins block cholesterol synthesis by suppressing HMG-CoA reductase^79^ and have been shown to be effective in NASH patients with dyslipidemia,^80, 81^ although other randomized trials with statins showed no significant improvement in liver histological features of NASH patients^82^ and raised concerns of statin-induced hepatotoxicity.^83^ Unlike statins, PCSK9 inhibitors reduce circulating cholesterol by inducing the uptake of cholesterol to the liver, potentially ameliorating NASH.^84^ In fact, PCSK9 knockout mice on high-cholesterol diets had increased hepatic cholesterol and fibrosing steatohepatitis,^21^ in line with our hepatocyte EHBP1–silenced NASH mice, where secreted PCSK9 is decreased. Although hepatocyte EHBP1 is beneficial in NASH, it may have negative effects on ASCVD due to its potential risk of inducing hypercholesterolemia. Therefore, targeting hepatocyte EHBP1 as a therapeutic strategy may not be ideal for treating NASH. Instead, the downstream mediator of EHBP1, *i.e.*, TAZ, can be a potential therapeutic target in NASH. Indeed, we have recently used siRNA conjugated to N-acetylgalactosamine (GalNAc),^85^ the first FDA-approved hepatocyte-target siRNA,^86^ to silence hepatocyte TAZ, and found that treatment of NASH mice with GalNAc-siTaz prevents and reverses fibrosis in NASH.^87^ In summary, the data in this report have revealed a novel hepatic cholesterol metabolism pathway by dissecting the interplay of the four cholesterol-associated GWAS loci in the context of NASH and provided new insight into cholesterol-based therapeutic intervention in NASH fibrosis.

### Limitations of Study

The *in vivo* causation experiments were conducted in an experimental model of NASH, and thus further work is needed to show the importance of the EHBP1-PCSK9 pathway in human NASH. Nonetheless, human relevance is supported by the following: (1) the loss-of-function *EHBP1* rs10496099 T>C variant is associated with augmented NASH in humans, which is consistent with EHBP1-silenced NASH mice; (2) the protein levels of EHBP1 and sortilin in hepatocytes are decreased in human NASH livers; (3) Sortilin and PCSK9 secretion are decreased in EHBP1-silenced primary human hepatocytes. In addition, whether EHBP1 regulates steatosis and inflammation independent of cholesterol metabolism in NASH, that is, via EHBP1-mediated endocytic trafficking of cell surface receptors that may participate in fatty acid uptake, remains unclear. Also, whether downstream mediators of EHBP1, including retromer subunits and sortilin, play roles in hepatic cholesterol metabolism and NASH progression awaits dedicated further studies. Our study shows that TNFα and PPARα modulate EHBP1 expression in primary hepatocytes, but whether they alter EHBP1-mediated cholesterol homeostasis *in vivo* is still unknown.

## STAR METHODS

Detailed methods are provided in the online version of this paper and include the following:

- KEY RESOURCES TABLE
- CONTACTS FOR REAGENT AND RESOURCE SHARING
- CELL CULTURE AND CELL ISOLATION
- AAV8 VIRAL VECTORS
- HISTOPATHOLOGICAL ANALYSIS FOR NASH MICE
- FILIPIN STAINING OF LIVER SECTIONS
- PLASMA AND CELL SUPERNATANT ANALYSIS
- IMMUNOBLOTTING
- CO-IMMUNOPRECIPITATION
- CHIP-SEQ ANALYSIS
- SNRNA-SEQ ANALYSIS
- RNA-SEQ ANALYSIS
- GENE ONTOLOGY (GO) ANALYSIS
- QUANTITATIVE RT-QPCR
- SIRNA-MEDIATED GENE SILENCING
- PHRODO RED LDL UPTAKE ASSAY
- IMMUNOFLUORESCENCE STAINING
- QUANTIFICATION AND STATISTICAL ANALYSIS

## SUPPLEMENTAL INFORMATION

Supplemental information includes seven Figures and six Tables.

### Key Resources Table

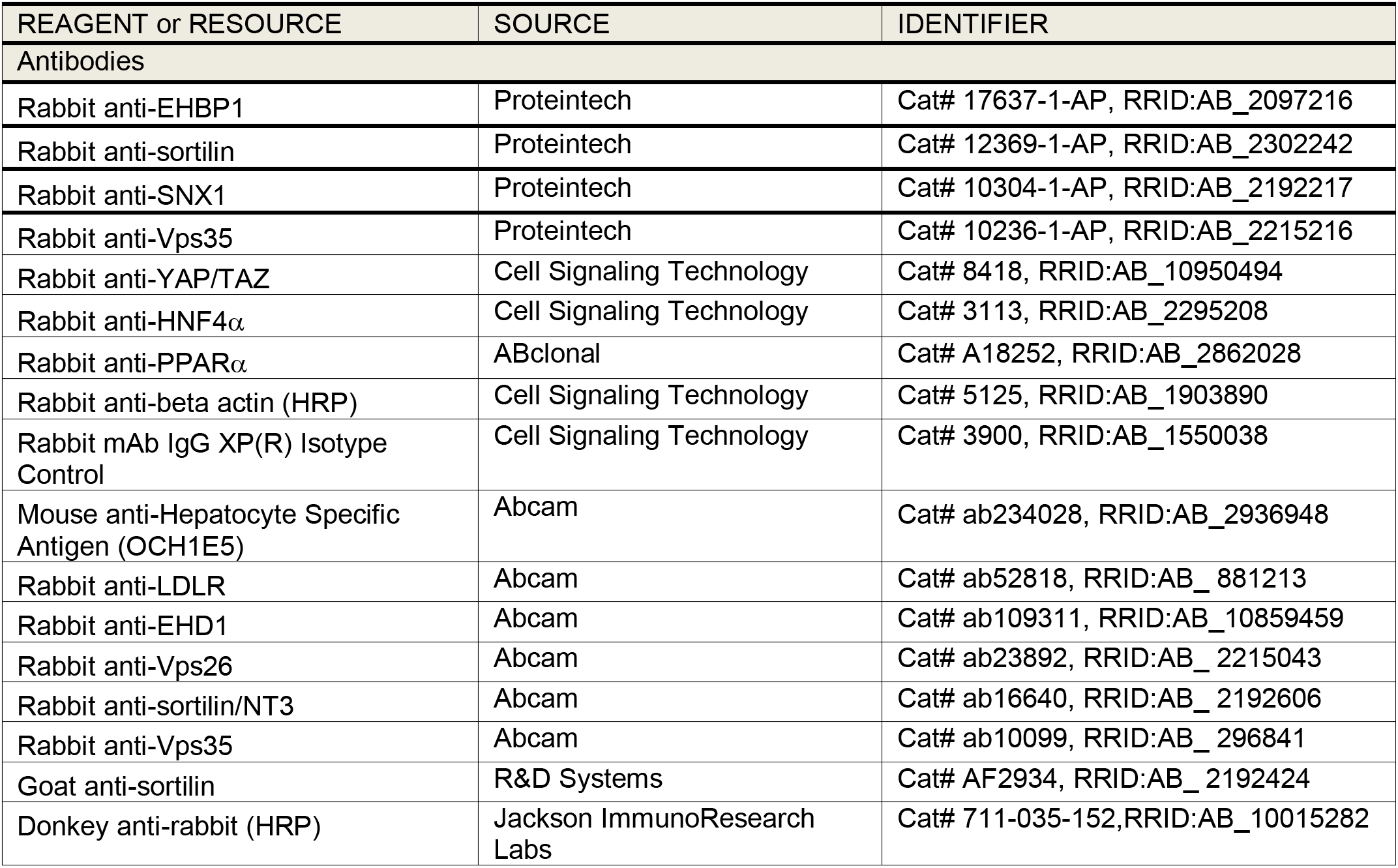
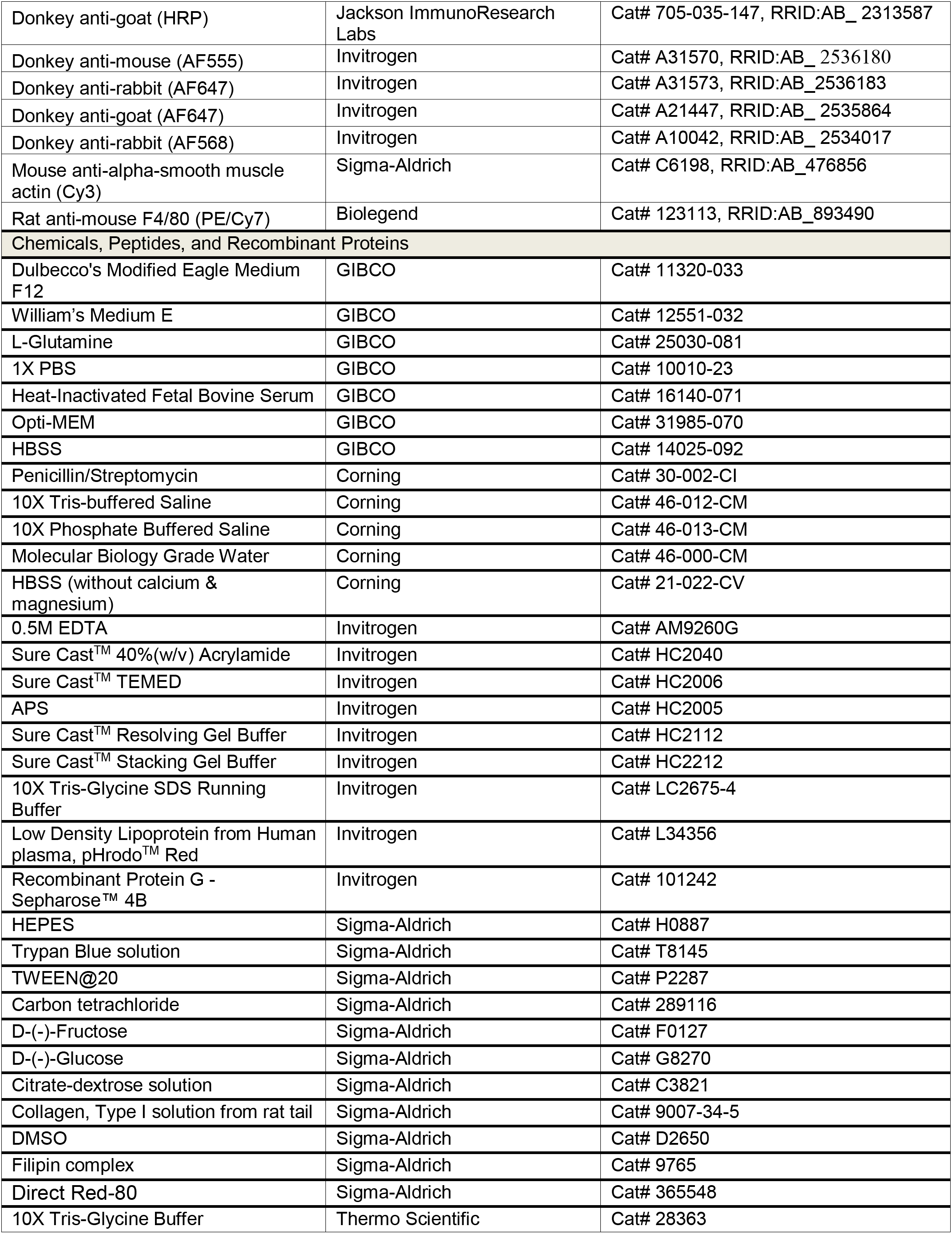
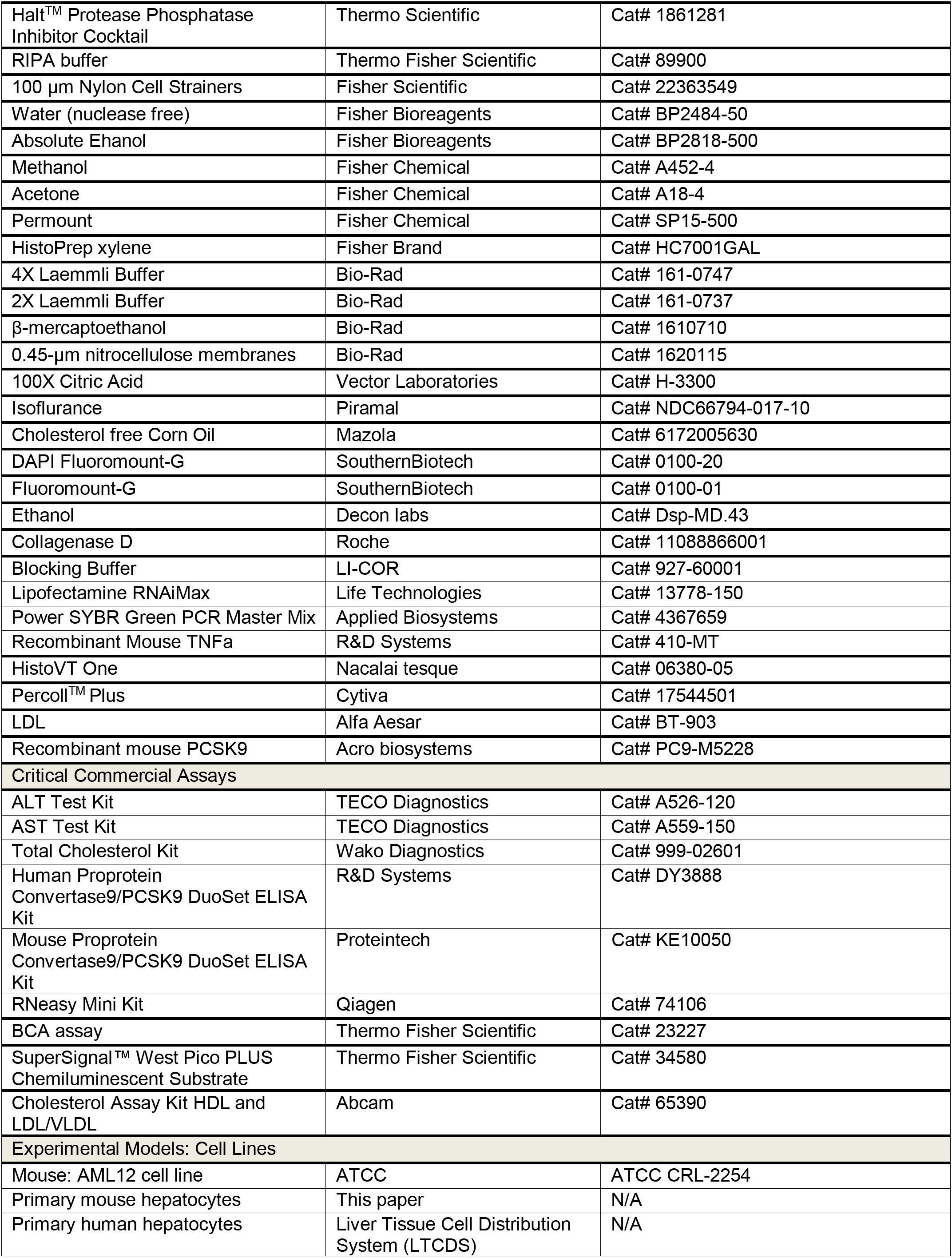
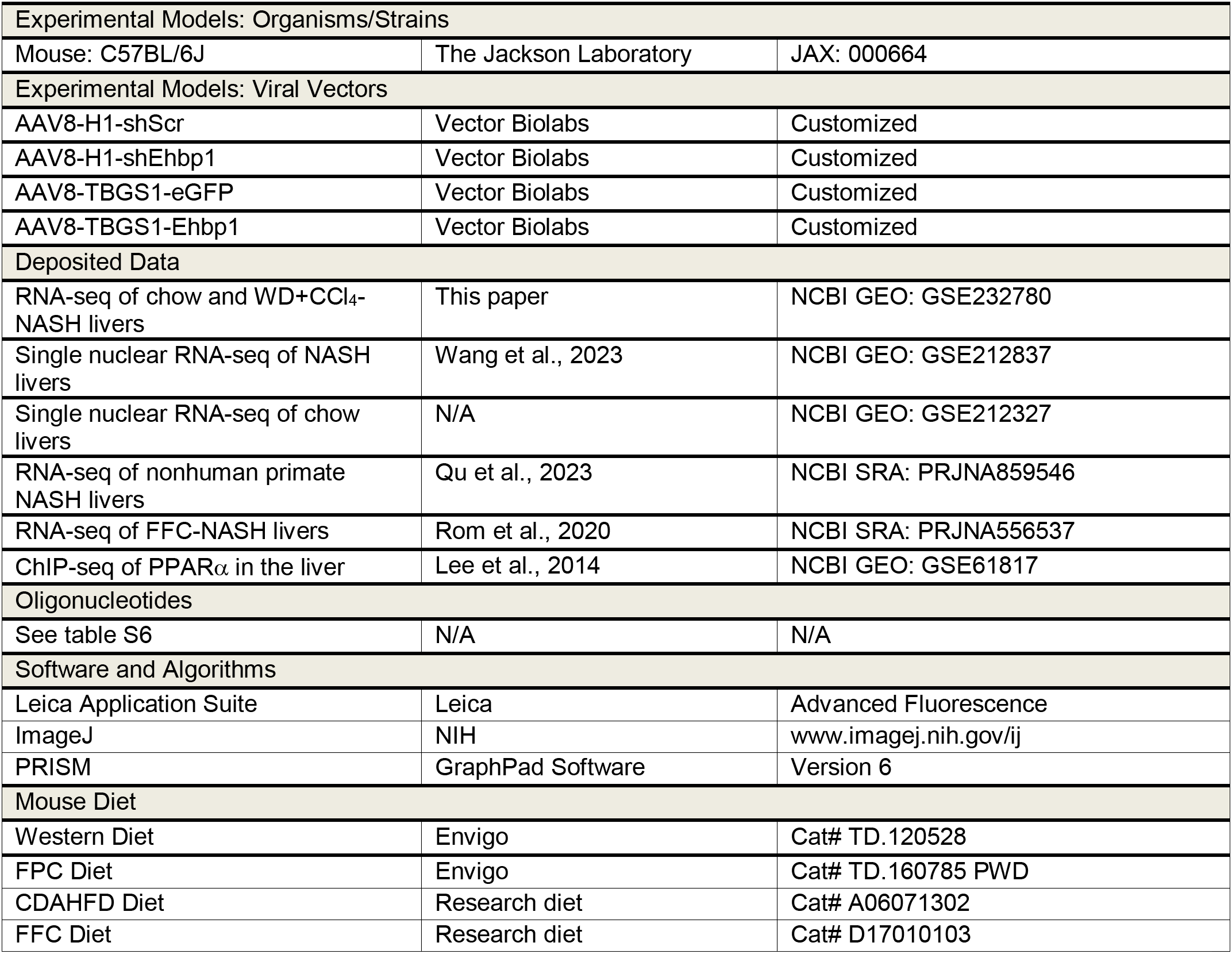

### Resource Availability

#### Lead Contact

Further information and requests for resources and reagents should be directed to and will be fulfilled by the lead contact, Dr. Bishuang Cai.

#### Materials Availability

All the materials generated in this study are available upon reasonable request to the lead contact. This study did not generate new unique reagents.

### Experimental Model and Subject Details

#### Experimental Animals

Male wild-type C57BL/6J (#000664, 7–10 weeks/old) mice were purchased from Jackson Laboratory (Bar Harbor, ME) and were allowed to adapt to housing in the animal facility for 1 week before the commencement of experiments. The mice were fed a NASH diet containing 21.1% fat, 41% sucrose, and 1.25% cholesterol by weight (Teklad diets, TD. 120528) and a high sugar solution (23.1 g/L d-fructose and 18.9 g/L d-glucose), with intra-peritoneally injected CCl_4_ at a dose of 0.2 µl (0.32 µg)/g of body weight once per week simultaneously with the diet administration for the times indicated in the figure legends, as previously described and validated in detail.^15^ AAV8-H1-shCon, AAV8-H1-shEhbp1, AAV8-TBGS1-eGFP, and AAV8-TBGS1-Ehbp1 viruses were delivered by tail vein injection at 2×10^11^ genome copies/mouse one week prior to NASH diet initiation. Five mice per cage were housed in standard cages at 22°C in a 12– 12 h light-dark cycle in a barrier facility. All procedures were performed according to protocols approved by the Animal Care and Use Committee of Icahn School of Medicine at Mount Sinai.

#### Human Liver and Primary Human Hepatocytes

For liver lysate analysis by immunoblotting and qPCR, deidentified human normal and NASH liver specimens as well as primary human hepatocytes were acquired from the Liver Tissue Cell Distribution System (LTCDS) at the University of Minnesota and the University of Pittsburgh. Phenotypic and pathological characterizations were conducted by physicians and pathologists associated with the LTCDS. To assess EHBP1 and sortilin staining in hepatocytes in human liver sections, deidentified human liver tissues were collected at time of surgery with informed consent at Mount Sinai Hospital, formalin-fixed, paraffin-embedded, and sectioned.^32^ Additional deidentified liver sections from individuals undergoing weight-loss surgery were selected from the MGH NAFLD Biorepository as we described previously.^88^ The diagnostic information is included in Table S5. All human studies were approved by the Icahn School of Medicine at Mount Sinai Institutional Review Board and were conducted in accordance with National Institutes of Health and institutional guidelines for human subject research.

#### Genotyping and Histologic Evaluation for the Hepatology Service Cohort

The Hepatology service cohort included 1367 patients with NAFLD unrelated patients of European descent who were enrolled consecutively at the Metabolic Liver Diseases outpatient service (Liver Clinic; n = 800) and bariatric surgery center (Bariatric Surgery; n = 567) at Fondazione IRCCS Cà Granda, Ospedale Maggiore Policlinico Milano (Milan, Italy) (Table S1).

Inclusion criteria were the availability of a liver biopsy specimen for suspected NASH or severe obesity, DNA samples, and clinical data. Individuals with excessive alcohol intake (men, >30 g/d; women, >20 g/d), viral and autoimmune hepatitis, or other causes of liver disease were excluded. The study conformed to the Declaration of Helsinki and was approved by the Institutional Review Board of the Fondazione Ca’ Granda IRCCS of Milan and relevant institutions. All participants provided written informed consent. NAFLD patients were genotyped for the *EHBP1* rs10496099 T>C variant. Genotyping was performed in duplicate using TaqMan 5’-nuclease assays (QuantStudio 3; Thermo Fisher, Waltham, MA). Results of rs10496099 T>C genetic frequency was compared with those obtained in non-Finnish European healthy individuals included in the 1000 Genome project3 (Table S2). Steatosis was divided into the following 4 categories based on the percentage of affected hepatocytes: 0, 0%–4%; 1, 5%–32%; 2, 33%–65%; and 3, 66%–100%. Disease activity was assessed according to the NAFLD activity score, with systematic evaluation of hepatocellular ballooning and necroinflammation. Fibrosis was also staged according to the recommendations of the NAFLD Clinical Research Network.^89^ The scoring of liver biopsy specimens was performed by independent pathologists unaware of patient status and genotype.^90, 91^ NASH was diagnosed in the presence of steatosis, lobular necroinflammation, and hepatocellular ballooning.

#### Gene Expression Analysis in Liver Biopsies from the Bariatric Surgery Cohort

*EHBP1* gene expression was measured in percutaneous liver biopsies of a subset of 125 severely obese patients from the Milan cohort (Bariatric surgery) whose clinical features are shown in Table S4. RNA was extracted from liver biopsies using RNeasy mini-kit (Qiagen Hulsterweg). RNA quality was assessed through Agilent 2100 Bioanalyzer and samples with RNA integrity numbers (RIN) greater than or equal to 7 were used for library preparation (Ribo reduction libraries). RNA sequencing was performed in paired-end mode with a read length of 150 nt using the Illumina HiSeq 4000 (Novogene, Hong Kong, China).

### Method Details

#### Cell Culture and Cell Isolation

Primary mouse hepatocytes were isolated from 7–10-week-old wild-type C57BL/6J mice as described previously.^92^ In brief, mice were euthanized with isoflurane, the abdomen was opened, and a cannula was inserted into the vena cava. The liver was perfused with Hanks’ balanced salt solution. Immediately upon appearance of white spots, the portal vein was cut and the liver was further perfused with collagenase D for digestion. After the digestion, the liver was placed in a petri dish containing DMEM and minced with forceps. The digested liver was passed through a 100 mm filter and centrifuged at 50 x g for 2 min. The supernatant fraction was removed, and the cell pellets were resuspended in 10 ml DMEM and 10 ml percoll (9 ml percoll + 1 ml 10 x PBS). The liver suspension was centrifuged at 200 x g for 10 min at 4°C. The supernatant was removed and the cell pellets (hepatocytes) were washed with 40 ml DMEM once. Washed hepatocytes were then plated in collagen-coated 24-well culture plates in DMEM/F12 containing 10% (vol/vol) heat-inactivated FBS and 1% pen-strep. For the analysis of sortilin-mediated PCSK9 secretion, primary hepatocytes were cultured in DMEM/F12 with 10% FBS for 5 h and then switched to William Medium E without FBS. AML12 mouse hepatocytes from ATCC (CRL-2254) were cultured in DMEM/F12 with 10% FBS and 1% pen-strep. All cells were cultured at 37°C and 5% CO_2_. The cells were harvested after treatment in Laemmli Sample Buffer (Bio-Rad) with 2-mercaptoethanol (Bio-Rad) for immunoblotting or in RNA lysis buffer for mRNA quantification.

#### AAV8 Viral Vectors

Adeno-associated virus subtype 8 (AAV8)-shRNA targeting murine Ehbp1 was made by annealing complementary oligonucleotides (5’-CACCAcacaaactgacgtcaagttaaCTCGAGTTAACTTGACGTCAGTTTGTG-3’) that were then ligated into the self-complementary (sc) AAV8-RSV-GFP-H1 vector, as described previously.^16^ The resultant constructs were amplified by Vector Biolabs, Malvern, PA. AAV8-TBGS1-eGFP and AAV8-TBGS1-Ehbp1 were purchased from Vector Biolabs.

#### Histopathological Analysis for NASH mice

Inflammatory cells in H&E-stained liver section images were quantified as the number of mononuclear cells per field (20x objective). Lipid droplets in H&E-stained liver section images were quantified as % lipid droplet area of total area by ImageJ. Liver fibrosis was detected with Sirius red staining. Briefly, rehydrated slides were stained for 1 h in saturated picric acid with 0.1% Sirius red (Direct Red-80; Sigma-Aldrich) followed by counterstain with 0.01% Fast Green (Sigma-Aldrich) for another 1 h. Liver fibrosis was quantified as % Sirius red^+^ area of total area by ImageJ. The same threshold settings were used for all analyses. For all analyses, we quantified 6–10 randomly chosen fields per section per mouse.

#### Filipin Staining of Liver Sections

Frozen liver sections were fixed in 4% paraformaldehyde for 10 min at room temperature, rinsed using glycine/PBS, and stained with 50 µg/ml filipin (Sigma-Aldrich) for 2 h in the dark. After washing with PBS, sections were mounted and viewed by fluorescence microscopy using a UV filter set (340–380 nm excitation, 40 nm dichroic, 430 nm long pass filter). As filipin fluorescence photobleaches very rapidly, care was taken to have the same UV exposure time before image collection for all samples.

#### Plasma and Cell Supernatant Analysis

Plasma alanine aminotransferase (ALT), aspartate aminotransferase (AST), and cholesterol levels in NASH mice were assayed using commercially available kits, as listed in the Reagents section. PCSK9 levels in mouse plasma, human serum, and culture supernatants from primary hepatocytes were assayed using PCSK9 ELISA kits (Proteintech; R&D Systems). LDL/VLDL cholesterol levels in human serum were assayed using the Cholesterol Assay kit (Abcam).

#### Immunoblotting

Liver protein lysates were extracted using RIPA buffer (Thermo Fisher Scientific), and protein concentration was measured using the BCA assay. Proteins were separated by electrophoresis on 4–20% Tris gels (Invitrogen) and transferred to 0.45 μm nitrocellulose membranes. The membranes were blocked for 1 h at room temperature in Tris-buffered saline/0.1% Tween 20 (TBST) containing 5% (wt/vol) nonfat milk and then incubated with the primary antibody in Intercept blocking buffer at 4°C overnight. The membranes were then incubated with the appropriate secondary antibody coupled to horseradish peroxidase, and proteins were detected by the SuperSignal™ West Pico PLUS Chemiluminescent Substrate (Thermo Fisher Scientific). Cultured cells were lysed in 2x Laemmli buffer containing 5% β-mercaptoethanol, heated at 100°C for 5 min, and then electrophoresed and immunoblotted as above.

#### Co-Immunoprecipitation (Co-IP)

Primary mouse hepatocytes or AML12 cells from 100-mm culture dishes were lysed in 2 ml ice-cold Co-IP buffer containing 25 mM Tris-HCL pH 7.5, 150 mM NaCl, 1.5 mM MgCl_2_, 1 mM EDTA, 0.5% NP-40, and protease inhibitor cocktail on ice for 30 min. Cell lysates were centrifuged at 10000 x g for 10 min at 4°C. The supernatant fraction was pre-cleaned using 30 µl protein G agarose beads (Invitrogen) for 2 h. Pre-cleaned supernatants were incubated with either rabbit anti-EHBP1 antibody or rabbit IgG (negative control) at 4°C overnight with gentle rotating, followed by incubation with per-washed protein G beads (50 µl) for 2 hours at 4°C. Beads were then collected by centrifugation and washed five times with 500 µl ice-cold Co-IP buffer. Proteins were eluted using 50 µl 2x Laemmli sample buffer and then subjected to immunoblot analysis.

#### ChIP-seq Analysis

PPARα ChIP-seq data were downloaded from NCBI Gene Expression Omnibus (GEO) with the accession number GSE61817. Reads were preprocessed by trim galore (v0.6.3) and aligned to the mm10 mouse genome using the bowtie2 (v2.3.4) with the default parameters. The aligned reads were sorted, and duplicates were removed with samtools (v1.0). The exported bam files were converted to a binary tiled file (tdf) and visualized using IGV (v2.7.2) software.

#### snRNA-seq Analysis

snRNA-seq datasets from two chow livers (GEO accession no. GSE212327) and two NASH livers (GEO accession no. GSE212837) were processed and annotated as we previously described.^32^ Expression of *Ehbp1* and *Ppara* in the hepatocyte population from chow and NASH livers was plotted using the DotPlot function in the Seurat (v4.2.0) package.

#### RNA-seq Analysis

Total liver RNA from 4 NASH and 4 chow mice was extracted using the RNeasy kit (Qiagen). RNA-seq library construction was performed at BGI group with a standard polyA-enrichment protocol. Sequencing was performed on an Illumina HiSeq 4000 Sequencer, and 100 bp paired end reads were obtained. For RNA-seq data processing, reads were aligned to the mouse genome mm10 using STAR (v2.7.6a) with the default settings. Transcript assembly and differential expression analyses were performed using Cufflinks (v2.2.1). Assembly of novel transcripts was not allowed (-G). Other parameters of Cufflinks were the default setting. The summed FPKM (fragments per kilobase per million mapped reads) of transcripts sharing each gene ID was calculated and exported by the Cuffdiff program. Differentially expressed genes (DEGs) were determined by two-sided T-test P-value <0.05 and fold-change >1.5. Volcano plots for gene expression by fold change versus P-value were generated using R.

#### Gene ontology (GO) analysis

Gene ontology (GO) analyses were performed for the DEGs using the DAVID gene ontology functional annotation tool (https://david.ncifcrf.gov/tools.jsp) with all *Mus musculus* genes as a reference list.

#### Quantitative RT-qPCR

Total RNA was extracted from liver tissue or cultured primary hepatocytes using the RNeasy kit (Qiagen). The quality and concentration of the RNA was assessed by absorbance at 260 and 280 nm using an EPOCH spectrophotometer (BioTek). cDNA was synthesized from 0.5–1 µg RNA using the High-capacity cDNA Reverse Transcription kit (Applied Biosystems). Quantitative RT-PCR was performed with a light cycler 480 II Real-time PCR system (Roche) using SYBR Green Master Mix (Bio-rad). The primer sequences were listed in Table S6.

#### siRNA-Mediated Gene Silencing

Negative control DsiRNA (# 51-01-14-04) and oligo-targeting siRNAs were purchased from IDT. The target sequence of mouse Ehbp1 siRNA (1) was AUUGGAUGGUGAAUUGG; the target sequence of mouse Ehbp1 siRNA (2) was ACACUGUUACUUAAAUCU; the target sequence of mouse Vps26a was GAAGUUUUCAGUAAGGUA; the target sequence of mouse Vps35 was AAGAUACCUGUUGAUACU; the target sequence of mouse SNX1 was GUCAGCAAAAUGACCAUC; the target sequence of mouse Ppara was GACCAAGUCACCUUGCUA; the target sequence of human EHBP1 was GGAGUAGACUUCUUAUCA. siRNAs were transfected into AML12 and primary hepatocytes using Lipofectamine RNAi MAX (Life Technologies) in 24-well plates following the manufacturer’s instructions. Briefly, 2 x 10^5^ cells at 30–40% confluence were incubated for ∼40 h in 0.5 ml of culture medium containing 1.5 µl Lipofectamine RNAiMAX and 20 pmol siRNA.

#### pHrodo Red LDL Uptake Assay

Primary mouse hepatocytes were seeded on cover glasses and transfected with siRNAs. Cells were then incubated with 5 μg/ml pHrodo Red LDL (Invitrogen) for 4 h. After washing three times with PBS, cells were mounted with DAPI-containing mounting solution. LDL uptake was detected by fluorescence microscopy and quantified from 6–10 randomly chosen fields per group as % LDL^+^ positive cells.

#### Immunofluorescence Staining

Primary mouse hepatocytes and AML12 cells were grown on cover glasses. Upon treatment, cells were fixed with 4% paraformaldehyde in PBS (pH 7.4) for 10 min and permeabilized with 0.1% Triton X-100 for 10 min at room temperature. Frozen liver sections were fixed in ice-cold acetone for 5 min and then incubated in 1 x Histo one buffer at 70°C for 1 h. Paraffin liver sections were de-paraffined in Xylene, rehydrated in graded ethanol (100%, 95%, 85%, and 70%), and subjected to antigen retrieval by placing in a pressure cooker for 10 min in citric acid buffer (Vector Laboratories). All samples were blocked with 5% donkey serum for 1 h and stained overnight at 4°C with primary antibodies, followed by incubation with secondary antibodies for 1 h at room temperature. Anti-fade solution was used for mounting cells on slides. Slides were viewed with a Leica TCS SP8 confocal fluorescence microscope or Zeiss fluorescence microscope.

#### Quantification and Statistical Analysis

All results are presented as mean ± SEM. Statistical significance was determined using GraphPad Prism software. P values were calculated using the Student’s t-test for normally distributed data and the Mann-Whitney rank sum test for non-normally distributed data. One-way ANOVA with Bonferroni post-test was used to analyze multiple groups with only one variable tested. Two-way ANOVA with Bonferroni post-test was used to analyze more than two groups with multiple variables tested. Correlation analysis were assessed by bivariate analysis (Pearson). All genetic analyses were performed under dominant model, according to the frequency distribution of the minor at-risk alleles among European non-Finnish healthy individuals included in the 1000 Genomes project. Analyses were performed by fitting data to multivariable models. In particular, nominal logistic regression models were fit to examine binary traits (steatosis ≥2, necroinflammation ≥1, fibrosis >1). When specified, confounding factors were included in the model.

#### Data and Code Availability

The RNA-seq data have been deposited at the Gene Expression Omnibus (GEO) with accession code: GSE232780. The deposited data are publicly available as of the date of publication. This paper analyzed existing, publicly available data. These accession numbers for the datasets were listed in the key resources table. This paper did not report original code. Any additional information required to reanalyze the data reported in this paper is available from the lead contact upon request.

## Acknowledgments

We thank the Liver Tissue Cell Distribution System at the University of Minnesota and the University of Pittsburgh for providing human normal and NASH liver tissues and primary human hepatocytes. This work was supported, in part, by NIH grants R00DK115778, R01DK134610, R01HL167107, R35GM147269, the Irma T. Hirschl/Monique Weill-Caulier Trust Research Award, and the PhRMA Foundation Research Starter Grant in Translational Medicine (to B.C.); R01DK128289, R01DK121154, and P30CA196521 (to S.L.F.); R21HD106263 (to X.H.); R01DK136016 and AGA2020-13-03 (to S.W.). R01DK134011 and R00HL150233 (to O.R.); R01HL167758 and R00HL145131 (to A.Y.J). Human liver samples were obtained from the Liver Tissue Cell Distribution System (University of Minnesota), funded by NIH contract HHSN276201200017C.

## Author Contributions

B.C. developed the study concept and experimental design; B.C., F.M., S.P., S.M., and S.H. conducted all the animal study procedures and the *in vitro* experiments; S.L.F. and D.B. provided advice related to the NASH study; X.W. helped make the shEhbp1 plasmid; X.H. and S.W. helped analyze RNA sequencing data; M.L. M.M., and E.P. genotyped the *EHBP1* rs10496099 T>C variant and performed the multivariate analysis for the Hepatology service cohort; P.D. supervised the experimental analysis in the Milan cohort; S.A. and O.R. helped analyze *EHBP1* and *PPARA* expression in nonhuman primate and FFC-induced NASH livers and provided *Ppara^fl/fl^* and *Ppara^HKO^* primary hepatocyte lysates; A.Y.J critically revised the manuscript; B.C. and F.M. interpreted the data and wrote the manuscript; and all coauthors participated in the manuscript editing.

---

**Figure S1 (Related to Figure 1).**
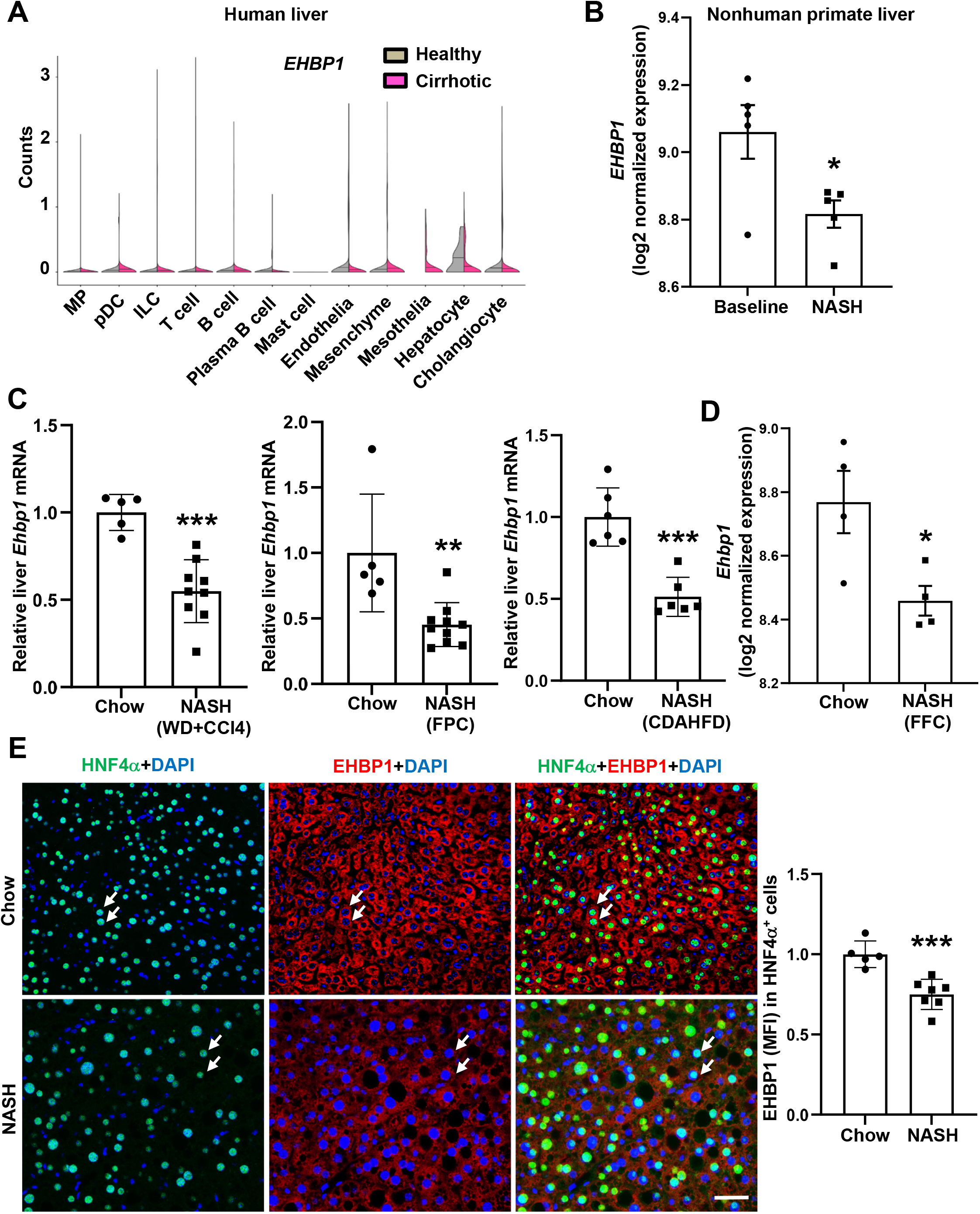
EHBP1 expression is reduced in human, nonhuman primate, and mouse NASH livers. (A) ScRNA-seq analysis for *EHBP1* expression in liver cells from healthy subjects or patients with cirrhotic livers. (B) RNA-seq analysis for *EHBP1* expression in nonhuman primate NASH livers (n = 5 nonhuman primates/group). (C) *Ehbp1* mRNA levels in WD+CCl_4_-, FPC-, CDAHFA-induced NASH livers (n = 5–10 mice/group). (D) RNA-seq analysis for *Ehbp1* expression in FFC-induced NASH livers (n = 4 mice/group). (E) EHBP1 (red) and HNF4α (green) immunofluorescence in sequential liver sections from 10-week NASH diet (WD+CCl_4_)-fed mice; quantification of EHBP1 (MFI) in HNF4α^+^ hepatocytes; DAPI counterstain for nuclei is shown and white arrows indicate HNF4α_+_ hepatocytes (n = 5–7 mice/group); Bar, 50 µm. Data are presented as mean ± SEM. *p < 0.05, **p < 0.01, ***p < 0.001.

**Figure S2 (Related to Figure 2).**
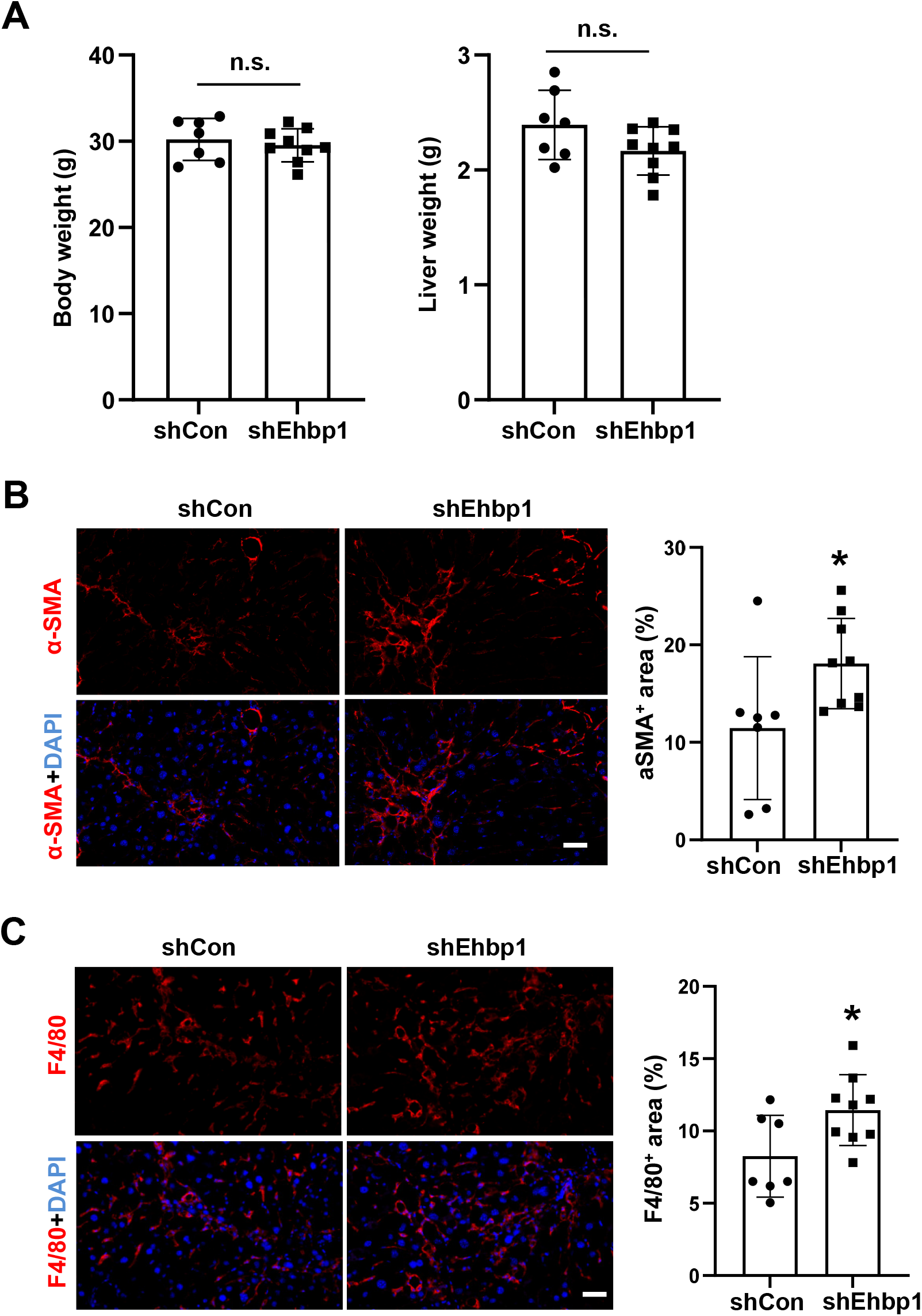
Fibrosis and inflammation are increased in hepatocyte EHBP1-silenced NASH livers. (A) Body and liver weights. (B) α-SMA immunofluorescence (red) and quantification; DAPI counterstain for nuclei is shown; Bar, 50 µm. (C) F4/80 immunofluorescence (red) and quantification; DAPI counterstain for nuclei is shown; Bar, 50 µm. Data are presented as mean ± SEM. *p < 0.05; n.s., no significance (n = 7–9 mice/group).

**Figure S3 (Related to Figure 3).**
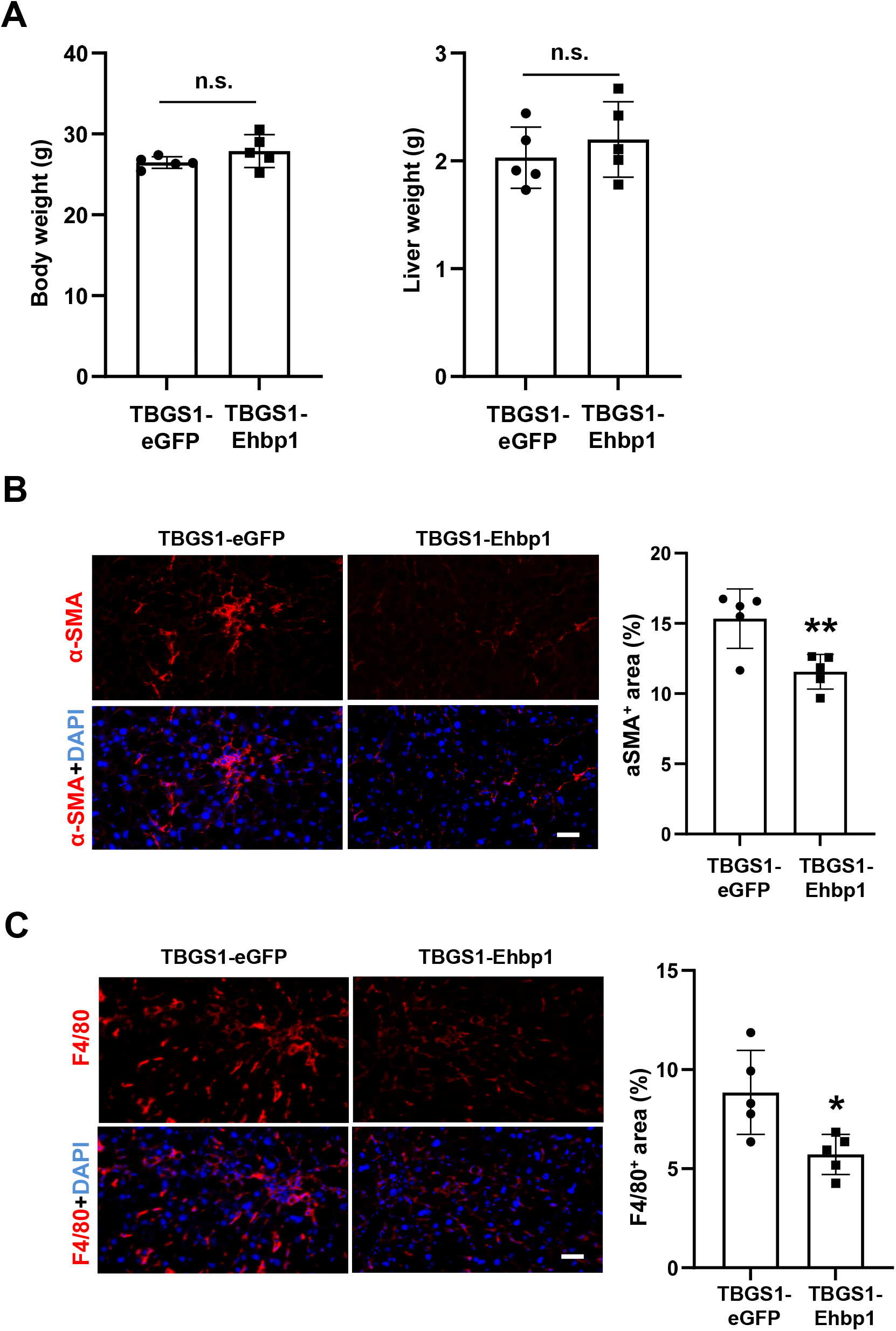
Fibrosis and inflammation are decreased in hepatocyte EHBP1-overexpressed NASH livers. (A) Body and liver weights. (B) α-SMA immunofluorescence (red) and quantification; DAPI counterstain for nuclei is shown; Bar, 50 µm. (C) F4/80 immunofluorescence (red) and quantification; DAPI counterstain for nuclei is shown; Bar, 50 µm. Data are presented as mean ± SEM. *p < 0.05, **p < 0.01; n.s., no significance (n = 5 mice/group).

**Figure S4 (Related to Figure 4).**
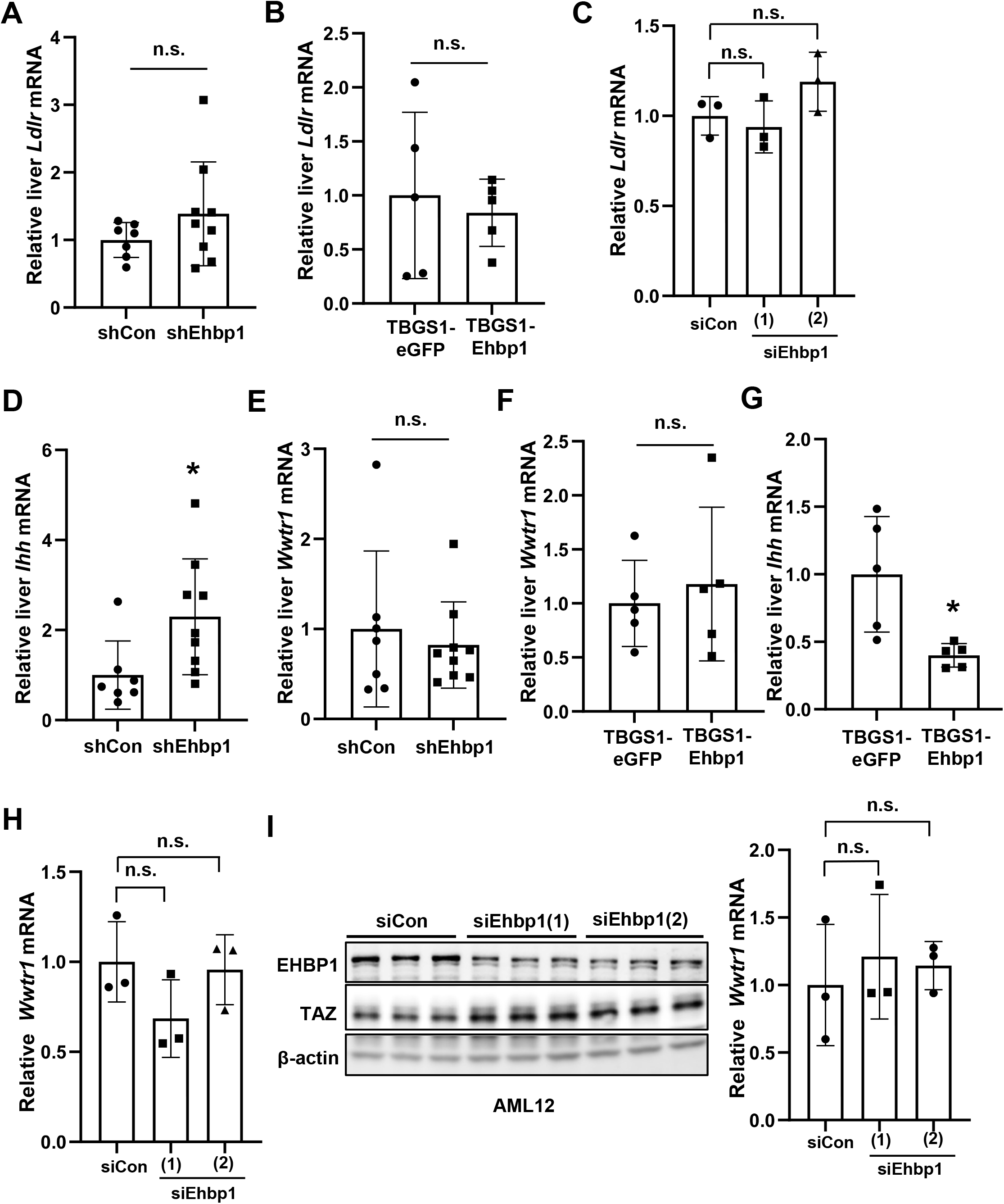
*Ldlr*, *Wwtr1*, *Ihh* expression in NASH livers and hepatocytes. (A) *Ldlr* mRNA level in AAV8-H1-shCon or AAV8-H1-shEhbp1-treated NASH livers (n = 7–9 mice/group). (B) *Ldlr* mRNA level in AAV8-TBGS1-eGFP or AAV8-TBGS1-Ehbp1-treated NASH livers (n = 5 mice/group). (C) *Ldlr* mRNA level in primary mouse hepatocytes treated with siCon or siEhbp1 (n = 3 biological replicates). (D) *Ihh* mRNA level in AAV8-H1-shCon or AAV8-H1-shEhbp1-treated NASH livers (n = 7–9 mice/group). (E) *Wwtr1* mRNA level in AAV8-H1-shCon or AAV8-H1-shEhbp1-treated NASH livers (n = 7–9 mice/group). (F) *Wwtr1* mRNA level in AAV8-TBGS1-eGFP or AAV8-TBGS1-Ehbp1-treated NASH livers (n = 5 mice/group). (G) *Ihh* mRNA level in AAV8-TBGS1-eGFP or AAV8-TBGS1-Ehbp1-treated NASH livers (n = 5 mice/group). (H) *Wwtr1* mRNA level in primary mouse hepatocytes treated with siCon or siEhbp1 (n = 3 biological replicates). (I) Immunoblots of EHBP1 and TAZ and *Wwtr1* mRNA level in AML12 cells treated with siCon or siEhbp1 (n = 3 biological replicates). Data are presented as mean ± SEM. *p < 0.05; n.s., no significance.

**Figure S5 (Related to Figure 5).**
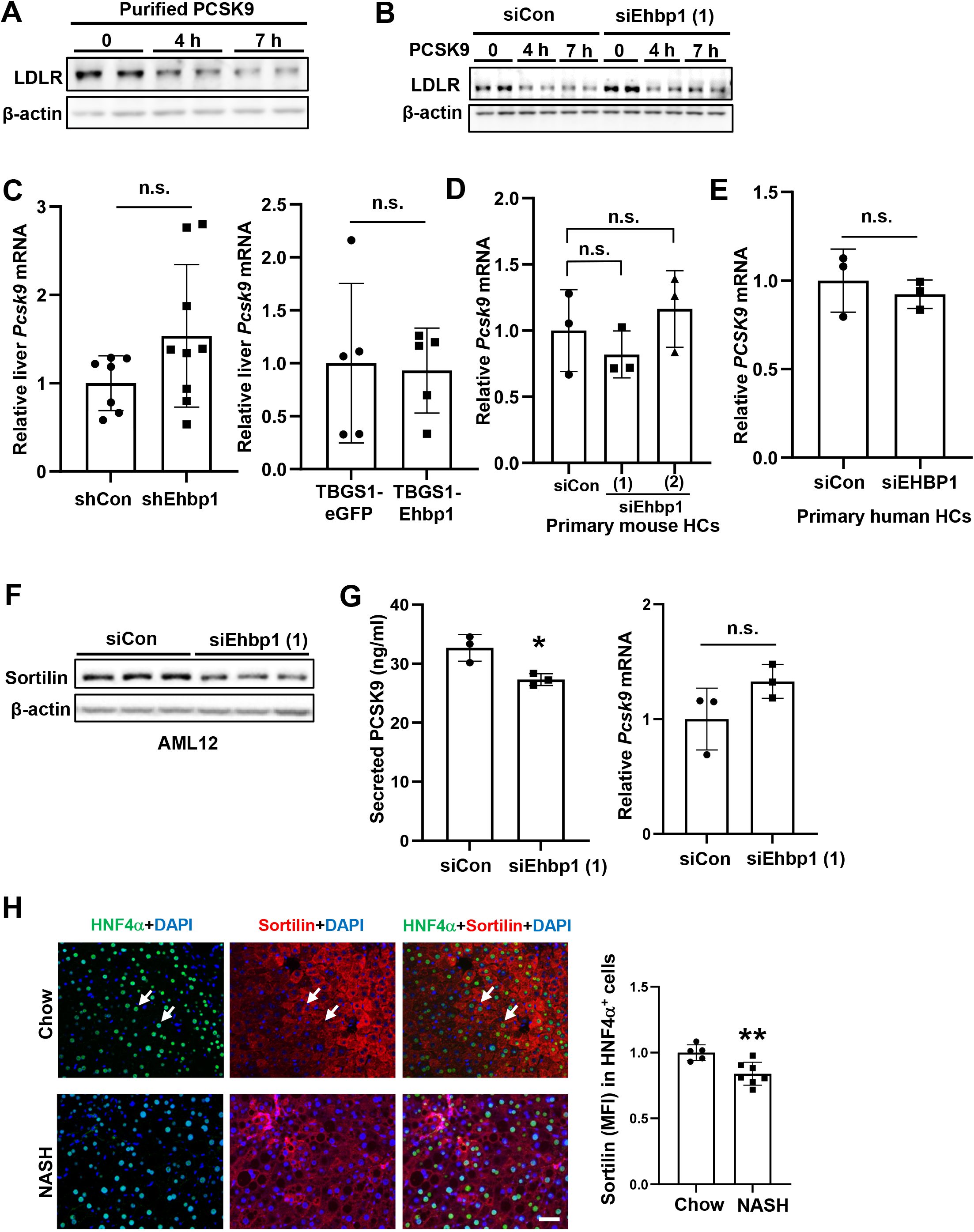
EHBP1 promotes sortilin-mediated PCSK9 secretion. (A) Immunoblot of LDLR in primary mouse hepatocytes incubated with 2.5 μg/ml PCSK9 for indicated time points. (B) Immunoblot of LDLR in siRNA-treated primary mouse hepatocytes followed by PCSK9 treatment. (C) *Pcsk9* mRNA levels in NASH livers (n = 5–9 mice/group). (D) *Pcsk9* mRNA level in primary mouse hepatocytes treated with siCon or siEhbp1 (n = 3 biological replicates). (E) *PCSK9* mRNA level in primary human hepatocytes treated with siCon or siEHBP1 (n = 3 biological replicates). (F) Immunoblot of sortilin in AML12 cells treated with siCon or siEhbp1. (G) Secreted PCSK9 level in culture media from and *Pcsk9* mRNA level in AML12 cells treated with siCon or siEhbp1 (n = 3 biological replicates). (H) Sortilin (red) and HNF4α (green) immunofluorescence in sequential liver sections from 10-week NASH diet (WD+CCl_4_)-fed mice; quantification of sortilin (MFI) in HNF4α^+^ hepatocytes; DAPI counterstain for nuclei is shown and white arrows indicate HNF4α hepatocytes (n = 5–7 mice/group); Bar, 50 µm. Data are presented as mean ± SEM. *p < 0.05, **p < 0.01; HC, hepatocyte; n.s., no significance.

**Figure S6 (Related to Figure 6).**
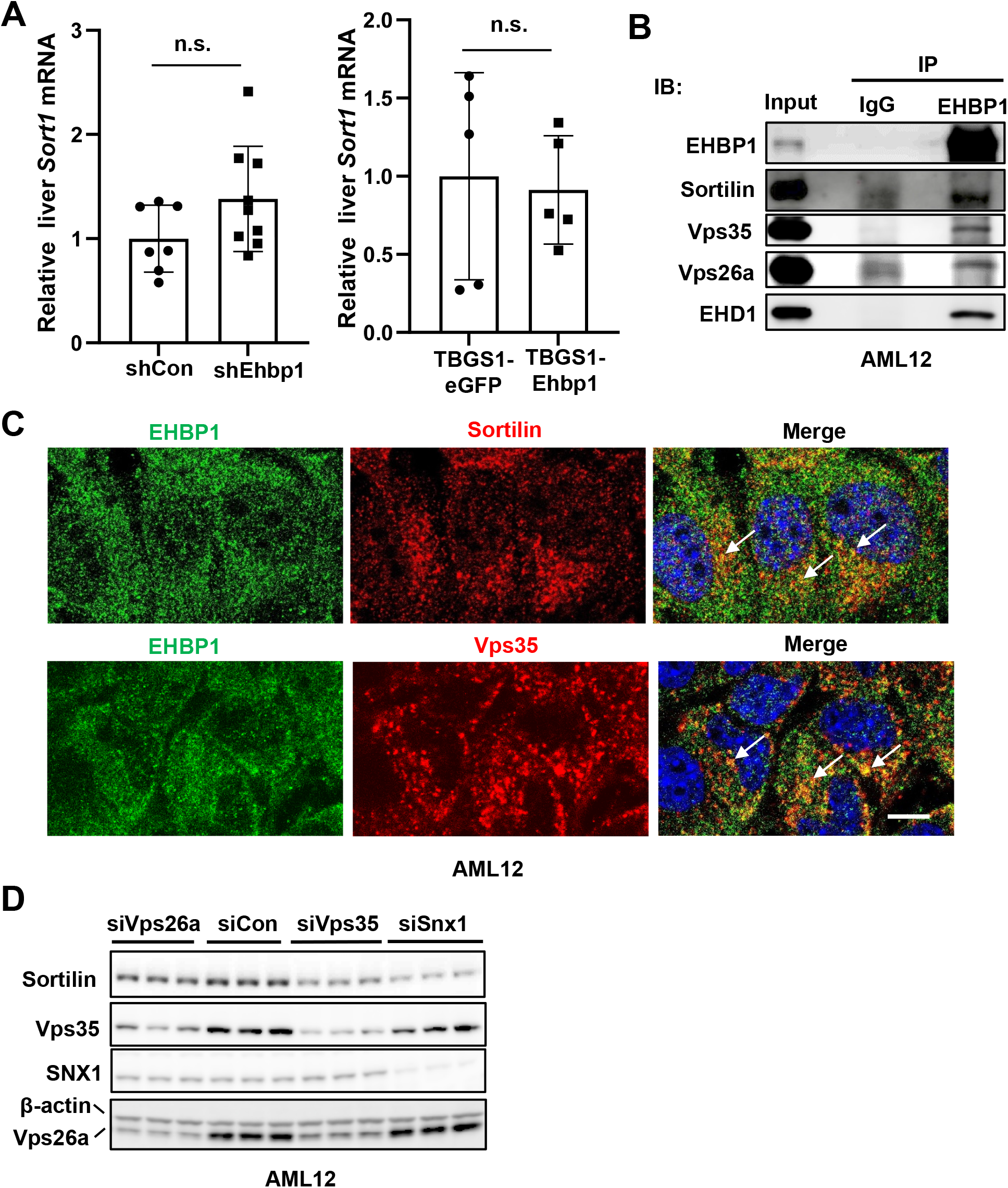
EHBP1 and sortilin form a complex with retromer subunits. (A) *Sort1* mRNA levels in NASH livers (n = 5–9 mice/group). (B) Co-immunoprecipitation of EHBP1, sortilin, Vps35, Vps26a, and EHD1 in AML12 cells. (C) Confocal imaging of EHBP1, sortilin, and Vps35 in AML12 cells; DAPI counterstain for nuclei is shown and white arrows indicate colocalization; Bar, 20 µm. (D) Immunoblots of sortilin, Vps35, SNX1, and Vps26a in AML12 cells treated with siRNAs. Data are presented as mean ± SEM. n.s., no significance.

**Figure S7 (Related to Figure 7).**
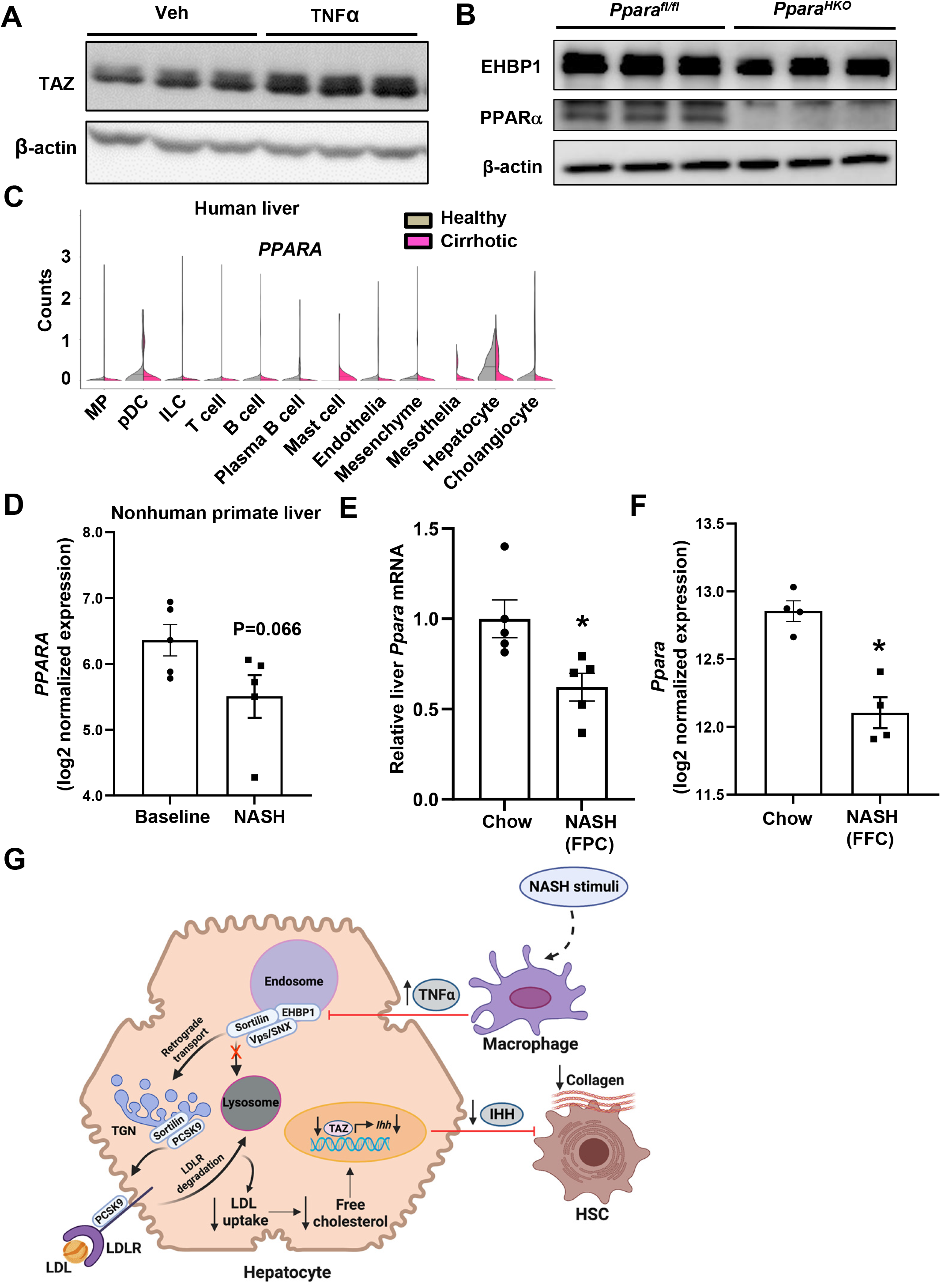
TNFα induces TAZ in hepatocytes and PPARα expression is reduced in human, nonhuman primate, and mouse NASH livers. (A) Immunoblot of TAZ in primary mouse hepatocytes that were incubated with 20 ng/ml TNFα for 24 h. (B) Immunoblots of EHBP1 and PPARα in primary mouse hepatocytes isolated from *Ppara^fl/fl^* and *Ppara^HKO^* (*Ppara^fl/fl^* Alb Cre*^+/-^*) mice. (C) ScRNA-seq analysis for *PPARA* expression in liver cells from healthy subjects or patients with cirrhotic livers. (D) RNA-seq analysis for *PPARA* expression in nonhuman primate NASH livers (n = 5 nonhuman primates/group). (E) *Ppara* mRNA level in FPC-induced NASH livers (n = 5 mice/group). (F) RNA-seq analysis for *Ppara* expression in FFC-induced NASH livers (n = 4 mice/group). (G) Summary scheme of EHBP1-mediated cholesterol metabolism. In homeostasis state, EHBP1 promotes sortilin-mediated PCSK9 secretion, leading to LDLR degradation and decreased LDL uptake into hepatocytes. The decreased uptake of LDL prevents free cholesterol accumulation, which in turn suppresses the stability of fibrogenic factor TAZ, decreases IHH expression, reduces HSC activation, and eliminates NASH fibrosis. However, as NASH progresses, macrophage-derived inflammatory cytokine TNFα abolishes the protective effect of EHBP1. HSC, hepatic stellate cell. Data are presented as mean ± SEM. *p < 0.05.

**Table S1.** Demographic, anthropometric, and clinical features of Hepatology service cohort (n = 1367), stratified according to the rs10496099 *EHBP1* T>C genotype.

|  | Hepatology Service cohort (n = 1367) | rs10496099 *EHBP1* |  | *P value |
| --- | --- | --- | --- | --- |
|  |  | TT (n = 280) | TC+CC (n = 1087) |  |
| Sex, M | 723 (52.88) | 142 (50.7) | 581 (53.4) | 0.36 |
| Age, years | 49.18±13.18 | 49.68 ±13.31 | 48.68±13.05 | 0.08 |
| BMI, kg/m<sup>2</sup> | 34.21±8.31 | 34.17±8.58 | 34.26±8.05 | 0.95 |
| HOMA-IR | 5.21±6.03 | 4.92 ±3.69 | 5.50±8.37 | 0.54 |
| Insulin, IU/ml | 21.08±1.50 | 20.52±1.98 | 21.64±1.03 | 0.70 |
| ALT, IU/l | 3.27 {3.46-3.6} | 3.49 {3.40-3.58} | 3.57 {3.53-3.62} | 0.18 |
| AST, IU/l | 3.29 {3.24 -3.62} | 3.27 {3.20 -3.33} | 3.32 {3.28-3.58} | 0.31 |

Values are reported as mean±SD, number (%) or median {IQR}, as appropriate. BMI: body mass index; HOMA-IR: homeostasis model assessment-estimated insulin resistance; ALT, alanine aminotransferase; AST, aspartate aminotransferase; Characteristics of participants were compared across the rs10496099 *EHBP1* T>C genotype using linear regression model (for continuous variables) or logistic regression model (for categorical characteristics).
\*Models were adjusted for sex, age, BMI, and presence of the rs738409 *PNPLA3* C>G variant.

**Table S2.** Genotype and allele frequencies of the rs10496099 *EHBP1* T>C variant in the Hepatology service cohort (n = 1367)

| rs10496099 | Alt Allele Frequency |
| --- | --- |
| Global | C=0.64 |
| European | C=0.63 |
| 1000G | C=0.60 |
| rs10496099 | NAFLD (n=1367) |
| TT (%) | 280 (20.48) |
| TC (%) | 679 (49.67) |
| CC (%) | 408 (29.84) |
| Total | 1367 |
| HWE | p<0.0001 |

**Table S3.** Association of *EHBP1* rs10496099 with liver damage in multivariable models in the Hepatology Service Cohort (n=1367)

| | Steatosis $\geq 2$ | | | Necroinflammation $\geq 1$ | | | Fibrosis $> 1$ | | |
| --- | --- | --- | --- | --- | --- | --- | --- | --- | --- |
|  | OR | 95% CI | P | OR | 95% CI | P | OR | 95% CI | P |
| Sex, M | 1.42 | 1.09-1.86 | 0.007 | 1.84 | 1.38-2.43 | <0.0001 | 1.79 | 1.30-2.45 | 0.0003 |
| Age, years | 1.00 | 0.99-1.01 | 0.93 | 1.03 | 1.02-1.04 | <0.0001 | 1.05 | 0.97-1.01 | <0.0001 |
| BMI, Kg/m<sup>2</sup> | 1.03 | 1.01-1.05 | <0.0001 | 1.03 | 1.02-1.05 | <0.0001 | 0.99 | 0.97-1.02 | 0.95 |
| PNPLA3 G allele, yes | 1.49 | 1.26-1.76 | <0.0001 | 1.35 | 1.05-1.75 | 0.01 | 1.64 | 1.34-2.00 | <0.0001 |
| EHBP1 C allele, yes | 1.42 | 1.06-1.91 | 0.03 | 1.35 | 1.00-1.83 | 0.04 | 1.51 | 1.02-2.20 | 0.03 |

OR: odd ratio; CI: confidence interval. Values were obtained at multivariate nominal regression adjusted for sex, age, BMI, and the *PNPLA3* rs738409 genetic variant.

**Table S4.**
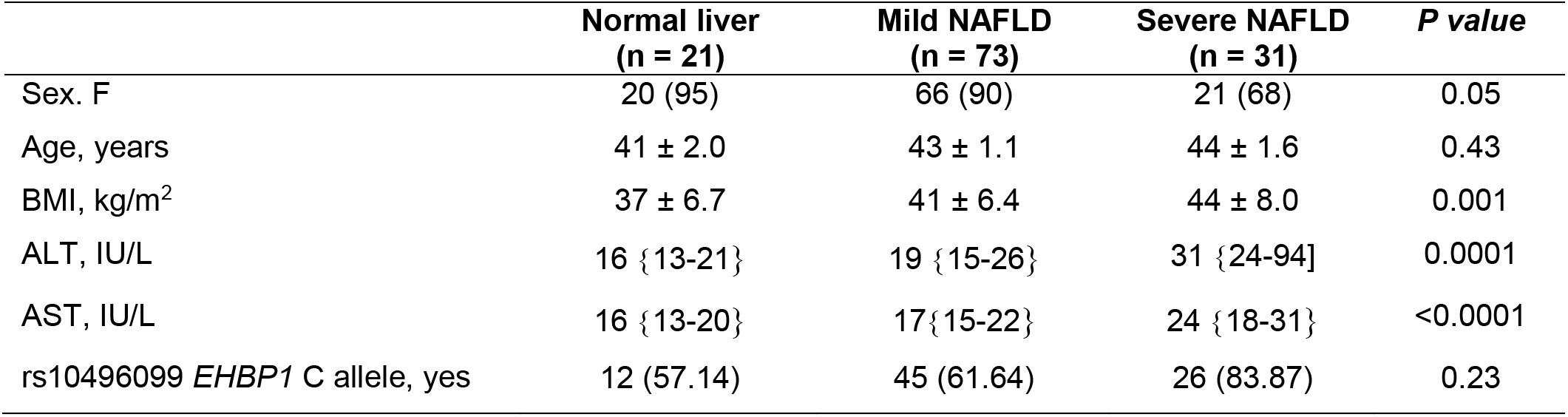
Demographic, anthropometric, and clinical features of 125 severely obese patients (Bariatric surgery), from whom RNA samples were available for RNA-seq analysis.

**Table S5.**
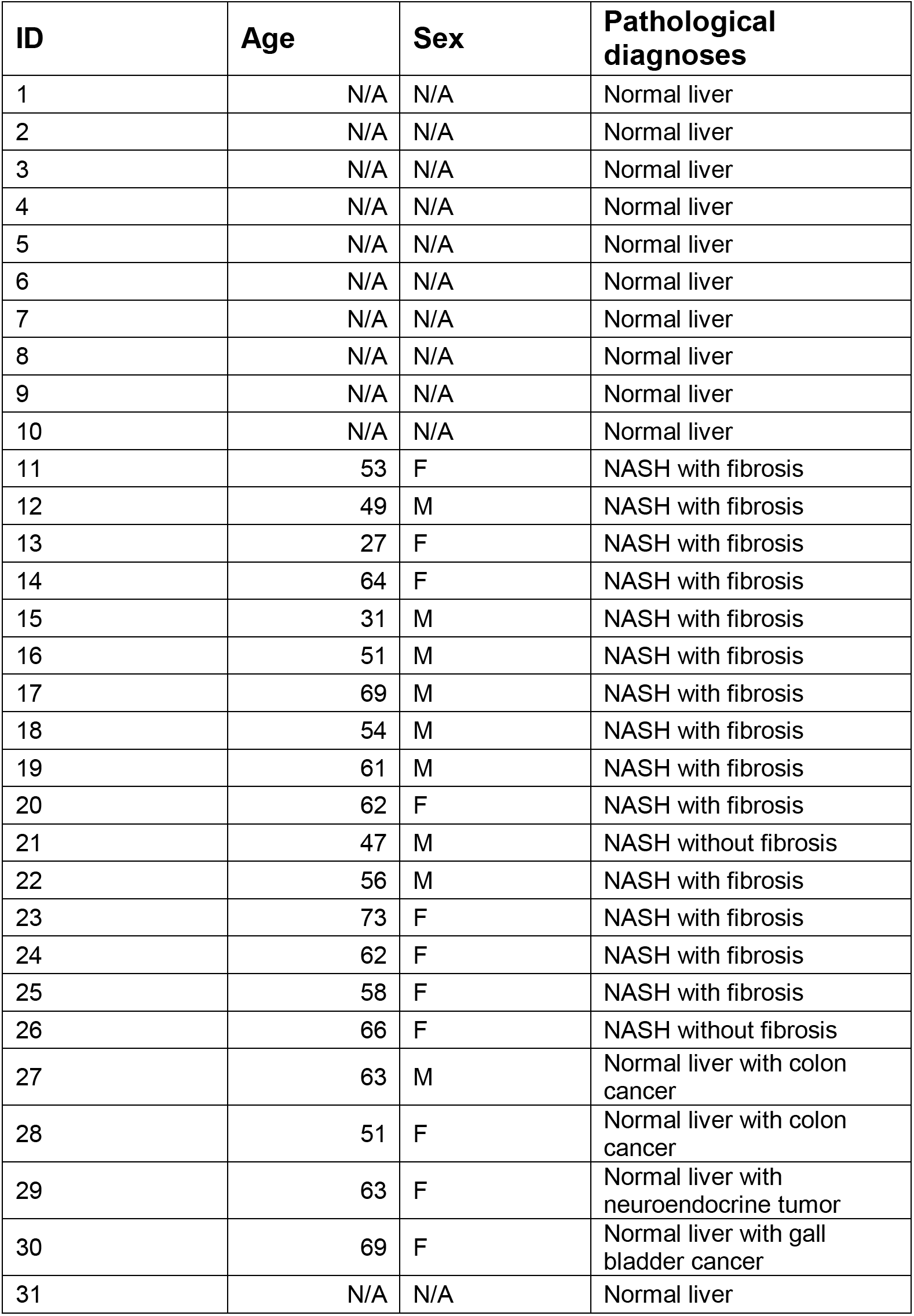
Pathology of human NASH liver samples for Figures 1A–1C and 5H.

| **ID** | **Age** | **Sex** | **Pathological diagnoses** |
| --- | --- | --- | --- |
| 1 | N/A | N/A | Normal liver |
| 2 | N/A | N/A | Normal liver |
| 3 | N/A | N/A | Normal liver |
| 4 | N/A | N/A | Normal liver |
| 5 | N/A | N/A | Normal liver |
| 6 | N/A | N/A | Normal liver |
| 7 | N/A | N/A | Normal liver |
| 8 | N/A | N/A | Normal liver |
| 9 | N/A | N/A | Normal liver |
| 10 | N/A | N/A | Normal liver |
| 11 | 53 | F | NASH with fibrosis |
| 12 | 49 | M | NASH with fibrosis |
| 13 | 27 | F | NASH with fibrosis |
| 14 | 64 | F | NASH with fibrosis |
| 15 | 31 | M | NASH with fibrosis |
| 16 | 51 | M | NASH with fibrosis |
| 17 | 69 | M | NASH with fibrosis |
| 18 | 54 | M | NASH with fibrosis |
| 19 | 61 | M | NASH with fibrosis |
| 20 | 62 | F | NASH with fibrosis |
| 21 | 47 | M | NASH without fibrosis |
| 22 | 56 | M | NASH with fibrosis |
| 23 | 73 | F | NASH with fibrosis |
| 24 | 62 | F | NASH with fibrosis |
| 25 | 58 | F | NASH with fibrosis |
| 26 | 66 | F | NASH without fibrosis |
| 27 | 63 | M | Normal liver with colon cancer |
| 28 | 51 | F | Normal liver with colon cancer |
| 29 | 63 | F | Normal liver with neuroendocrine tumor |
| 30 | 69 | F | Normal liver with gall bladder cancer |
| 31 | N/A | N/A | Normal liver |

**Table S6.**
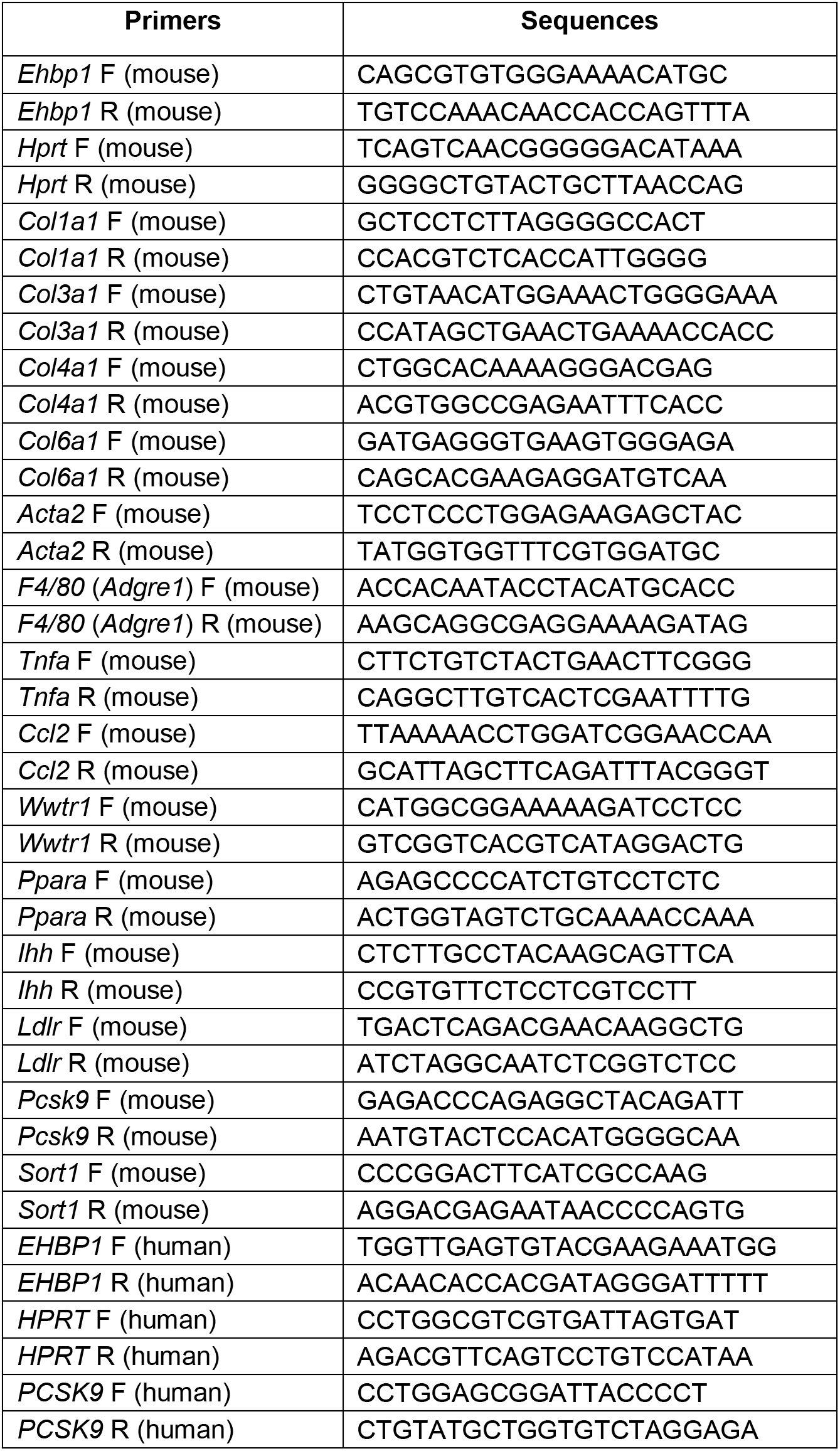
(Related to multiple figures). Primers used for QPCR.

